# Transcript isoform sequencing reveals widespread promoter-proximal transcriptional termination

**DOI:** 10.1101/805853

**Authors:** Ryan Ard, Quentin Thomas, Bingnan Li, Jingwen Wang, Vicent Pelechano, Sebastian Marquardt

## Abstract

Higher organisms achieve optimal gene expression by tightly regulating the transcriptional activity of RNA Polymerase II (RNAPII) along DNA sequences of genes^1^. RNAPII density across genomes is typically highest where two key choices for transcription occur: near transcription start sites (TSSs) and polyadenylation sites (PASs) at the beginning and end of genes, respectively^2,3^. Alternative TSSs and PASs amplify the number of transcript isoforms from genes^4^, but how alternative TSSs connect to variable PASs is unresolved from common transcriptomics methods. Here, we define TSS/PAS pairs for individual transcripts in *Arabidopsis thaliana* using an improved Transcript Isoform sequencing (TIF-seq) protocol and find on average over four different isoforms corresponding to variable TSS/PAS pairs per expressed gene. While intragenic initiation represents a large source of regulated isoform diversity, we discover that ∼ 14% of expressed genes generate relatively unstable short promoter-proximal RNAs (sppRNAs) from nascent transcript cleavage and polyadenylation shortly after initiation. The location of sppRNAs coincides with increased RNAPII density, indicating these large pools of promoter-stalled RNAPII across genomes are often engaged in transcriptional termination. RNAPII elongation factors progress transcription beyond sites of sppRNA formation, demonstrating RNAPII density near promoters represents a checkpoint for early transcriptional termination that governs full-length gene isoform expression.

---

Alternative TSS and PAS selection allows organisms to expand RNA isoform diversity from individual genes. Such alternative isoforms may encode proteins with differing activities characteristic of disease^5,6^ or produce functional long non-coding RNA (lncRNA) molecules^7^. Since commonly used transcriptomics methods focus on detecting either the TSS or the PAS of transcription unit (TU) boundaries, the molecular mechanisms connecting alternative TSSs to variable PASs yielding different gene isoforms are poorly understood. To overcome this challenge, we mapped corresponding TSS/PAS pairs for individual RNA molecules genome-wide in *Arabidopsis thaliana* seedlings using an improved TIF-seq protocol (see Methods; **Fig. S1-3**)^4^. TIF-seq detected TU boundaries for thousands of putative novel gene isoforms, such as those visible at the *GT-2 LIKE 1* (*GTL1*) gene (**Fig. 1a**). TIF-seq validated previously characterized alternative gene isoforms in *Arabidopsis* that result from alterative TSSs or PASs (**Fig. S3d-e**), as well as antisense lncRNA variants (**Fig. S3f**). We combined TUs with 5′ and 3′ end sites co-occurring within 20 nt windows into clusters, and eliminated those with 5’ and 3’ mispriming, yielding 50 thousand unique “TIF-clusters” (see Methods; **Table S1**). These analyses estimate an average of 4.3 distinct isoforms corresponding to alternative TSS/PAS pairs per expressed gene (**Fig. S3c**). While most TSS/PAS pairs map annotated gene boundaries (**Fig. 1b-d**), considerable variability exists in 5’- and 3’-untranslated region (UTR) length (**Fig. 1e**). These variations might impact RNA stability, targeting, and/or translation^8^. Intragenic initiation that terminates at canonical gene ends also represents a common origin of plant isoform diversity (**Fig. 1d**). Intragenic TSSs are suppressed by a chromatin-based mechanism involving the conserved RNAPII-associated histone chaperone complex Facilitates Chromatin Transcription (FACT), consisting of SPT16 and SSRP1^9^. TIF-seq in mutants of both *Arabidopsis* FACT subunits revealed that intragenic TSSs produce alternative gene isoforms that terminate at gene ends (**Fig. 1f; Fig. S4**). Thus, intragenic initiation regulated by FACT connects with canonical PASs in *Arabidopsis* to yield alternative gene isoforms. Overall, our TIF-seq data reveal thousands of alternative TUs encoded by the *Arabidopsis* genome.

**Fig. 1.**
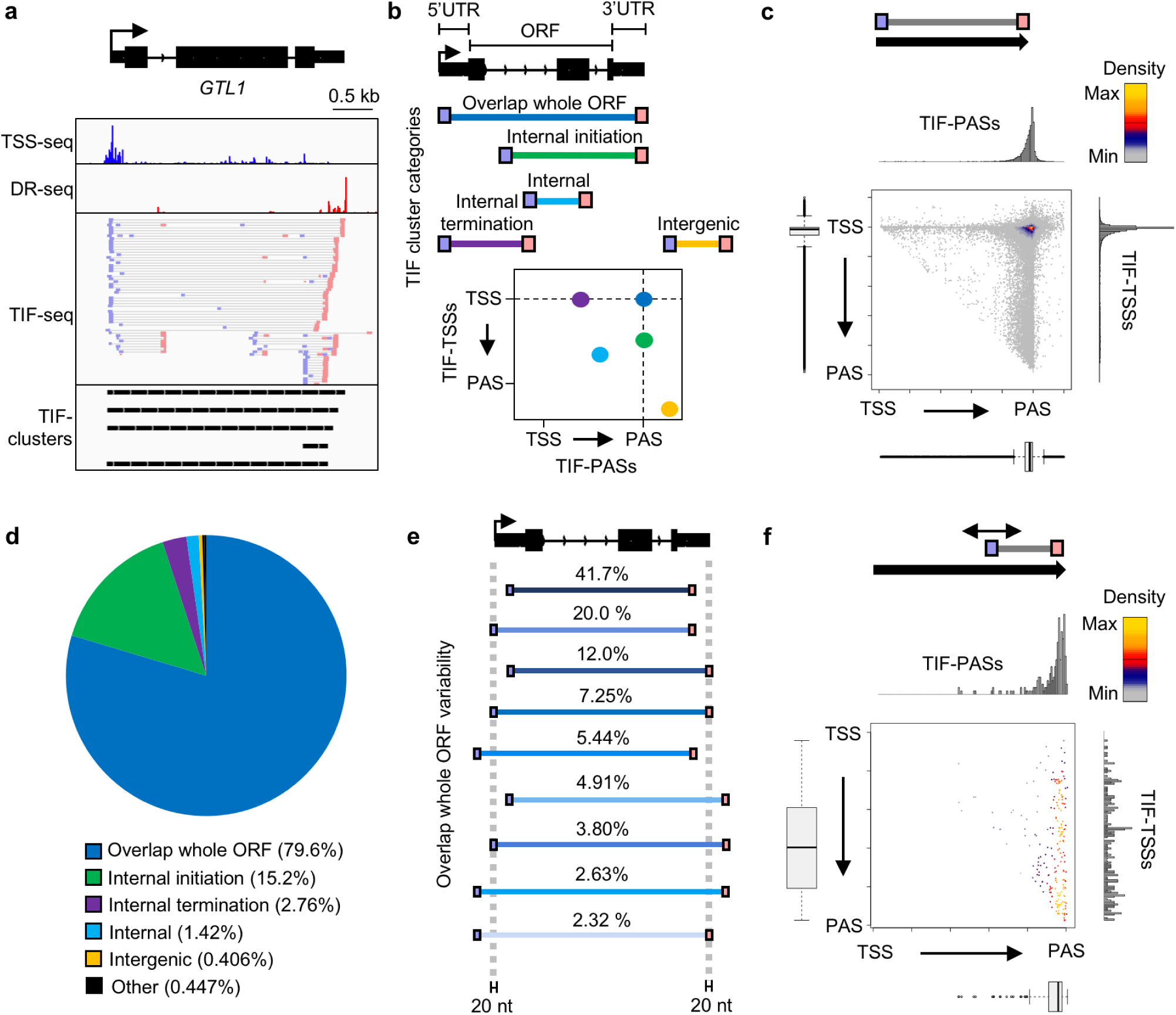
Mapping TU boundaries in *Arabidopsis* by TIF-seq. (a) Genome browser screenshot of TSSs (TSS-seq)^9^, PASs (Direct RNA-seq; DR-seq)^30^, TSS/PAS pairs (TIF-seq), and TIF-clusters at the *GTL1* gene. (b) Schematic of TIF-TSS/PAS pairs representing different TIF Cluster Categories and 2D illustration of these positions with respect to genome annotations. (c) Scatterplot of TIF-cluster TSS/PAS pairs in wild type. Histograms and bar plots display TIF-TSSs/-PASs distribution normalized to gene length from annotated TSS to PAS. (d) Piechart of TIF Cluster Categories proportions present in wild type. (e) Schematic representation and percentage of subcategories for TIF-clusters that overlap whole ORFs. (f) Scatterplot of TSS/PAS pair positions for TIF-clusters originating from intragenic FACT-repressed TSSs in *fact* mutants (see Methods).

RNAPII transcription is tightly linked to RNA degradation by the nuclear exosome complex, which acts as a surveillance mechanism to eliminate defective or superfluous transcripts^10^. The detection of many RNA species that are produced in wild type yet rapidly degraded (“cryptic RNA”) is facilitated by analyses in nuclear exosome mutants. *Arabidopsis* mutants lacking the RNA helicase HUA ENHANCER 2 (HEN2) display defective exosome activity^11^. We performed TIF-seq in the *hen2-2* mutant to map cryptic TUs in *Arabidopsis* (**Fig. S5**). An increased proportion of intergenic and antisense TUs were observed by TIF-seq in *hen2-2* (**Fig. S5**), exceeding previous microarray analyses^11^. Interestingly, we discovered a high proportion of prematurely terminated transcripts enriched near promoter regions (**Fig. 2a**). Size distribution plots show that short TUs accumulate in *hen2-2* (**Fig. 2b**). These short TUs (<200 nts) mostly terminate close to their respective promoter TSSs (**Fig. 2c**). We defined this novel transcript class as short promoter-proximal RNAs (sppRNAs), with a median length of ∼ 93 nts (**Fig. S5e**). In total, we detected sppRNAs at ∼ 14% of expressed genes (**Table S1**). Interestingly, genes with sppRNAs show elevated gene body RNAPII transcription (**Fig. S6**), indicating sppRNAs are detected at genes with generally higher RNAPII transcriptional activity. Collectively, TIF-seq revealed sppRNAs as a prevalent feature of plant gene expression.

**Fig. 2.**
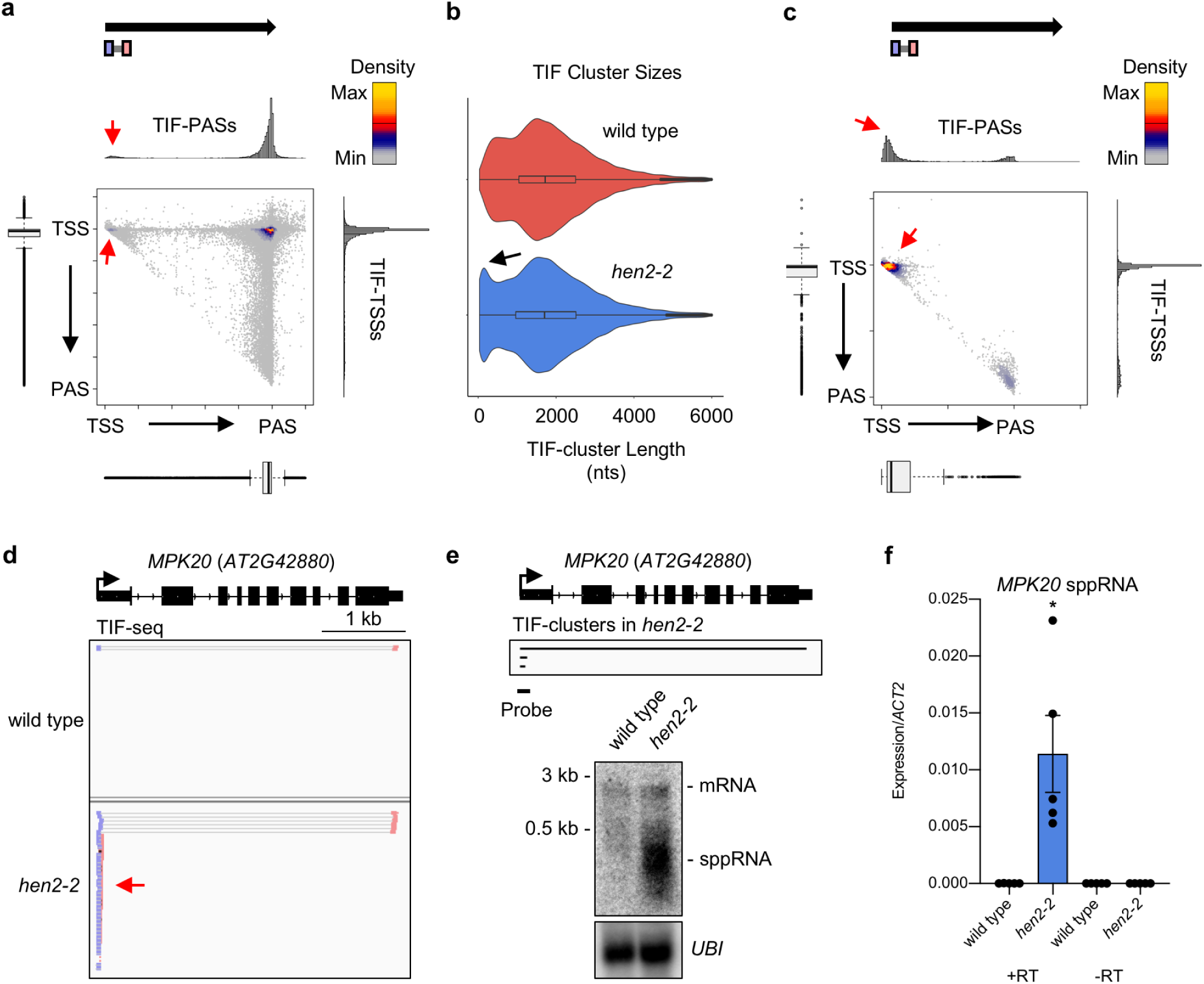
TIF-seq in *hen2-2* reveals promoter-proximal termination. (a) Scatterplot of TIF-cluster TSS/PAS pairs in *hen2-2*. Histograms and bar plots display TIF-TSSs/-PASs distribution normalized to gene length from annotated TSS to PAS. Red arrows indicate TSS/PAS pairs within initiation regions. (b) TIF-cluster sizes in wild type (n=41,190) and *hen2-2* (n=48,225). Black arrow indicates enrichment of short TUs in *hen2-2*. (c) Scatterplot of TIF-clusters <200 nt in *hen2-2*. (d) Genome browser screenshot of full-length *MPK20* mRNA and cryptic *hen2-2*-specific sppRNAs. (e) Northern blotting against *MPK20* sppRNAs in wild type and *hen2-2. UBI* is loading control. TIF-clusters in *hen2-2* and northern probe position indicated. (f) RT-qPCR for *MPK20* sppRNA levels in wild type and *hen2-2.* With reverse transcriptase (+RT) and without (-RT). Black dots indicate individual data-points. Error bars represent SEM resulting from five independent experiments. Asterisk denotes p<0.05 between wild type and *hen2-2* by Student’s t-test.

We validated sppRNAs by alternative methods at six genes with varying steady-state mRNA levels (**Fig. 2d; Fig. S7**). TIF-seq predicts prominent *hen2-2*-specific sppRNAs at the *MAP KINASE 20* (*MPK20*) locus (**Fig. 2d**). Consistent with alternative sppRNA PASs, we detected a broad size distribution for *MPK20* sppRNAs in *hen2-2* by northern blotting (**Fig. 2e**). Further validation was achieved by more sensitive RT-qPCR, which identified low *MPK20* sppRNA levels in wild type that significantly accumulated in *hen2-2* (**Fig. 2f**). While sppRNAs at additional genes also generally increased in *hen2-*2, low sppRNA levels were detected above background in wild type (**Fig. S8**). Interestingly, sppRNA accumulation in *hen2-2* or a point mutation in the conserved Ribosomal RNA Processing 4 (RRP4) subunit of the nuclear exosome^12^ did not significantly alter mRNA levels for tested genes (**Fig. S8-10**), suggesting sppRNA stabilization does not significantly impact cognate mRNA levels. Together, these results show that premature termination near *Arabidopsis* promoters generates relatively short-lived capped and polyadenylated transcripts that are targeted by the nuclear exosome RNA degradation machinery.

Transcription events initiating at the same TSS can form either full-length mRNAs or sppRNAs, which could represent a mechanism to adjust gene expression by selective transcriptional termination^3^. Temperature can affect the efficiency of transcriptional termination^13^. Moreover, GO-term analyses reveal that a significant number of genes with sppRNAs are cold-regulated (**Table S2**). To test the possible connection of temperature change and gene regulation by sppRNA formation, we performed TIF-seq in wild type and *hen2-2* seedlings following 3 hours of cold treatment (**Fig. S11-12)**, which is sufficient to trigger coding and lncRNA transcriptional responses that promote cold acclimation^14^. We compared wild type TSS-seq data that detects the mRNA of sppRNA genes in both conditions to identify regulation of mRNA formation at the expense of sppRNAs following cold (**Fig. S13a**). mRNA increases at the expense of sppRNAs for 38 sppRNA genes following cold treatment (**Fig. S13b**), suggesting selective termination may contribute to transcriptome adaptations to cold. However, most genes with sppRNA show no evidence for gene regulation by selective termination and an equivalent fraction of mRNA without sppRNAs are cold-induced (**Fig. S13b**). Interestingly, we detect sppRNAs at many cold-induced genes following cold treatment (**Fig. S13c-d**), consistent with sppRNA formation linked to greater RNAPII activity (**Fig. S6**). Thus, promoter-proximal termination is associated with plant gene expression across temperatures and may contribute to temperature-dependent gene regulation.

We predicted that sppRNAs result from promoter-proximal RNAPII regulation. To examine nascent RNAPII activity, we plotted plant Native Elongating Transcript sequencing (pNET-seq) data for genes with detectable sppRNAs^15^. The end positions of sppRNAs (i.e. PAS) map within peaks of RNAPII occupancy (**Fig. 3a-b**). This pattern is also observed for pNET-seq using a different RNAPII antibody (**Fig. S14**)^15^. In fact, RNAPII stalling was specifically enriched near promoters for genes with sppRNAs when compared against a control set of genes without detectable sppRNAs but otherwise equal gene body RNAPII activity (**Fig. 3c**). Overall, significantly increased RNAPII stalling shortly after initiation correlates strongly with promoter-proximal cleavage and polyadenylation generating sppRNAs (**Fig. S15**).

**Fig. 3.**
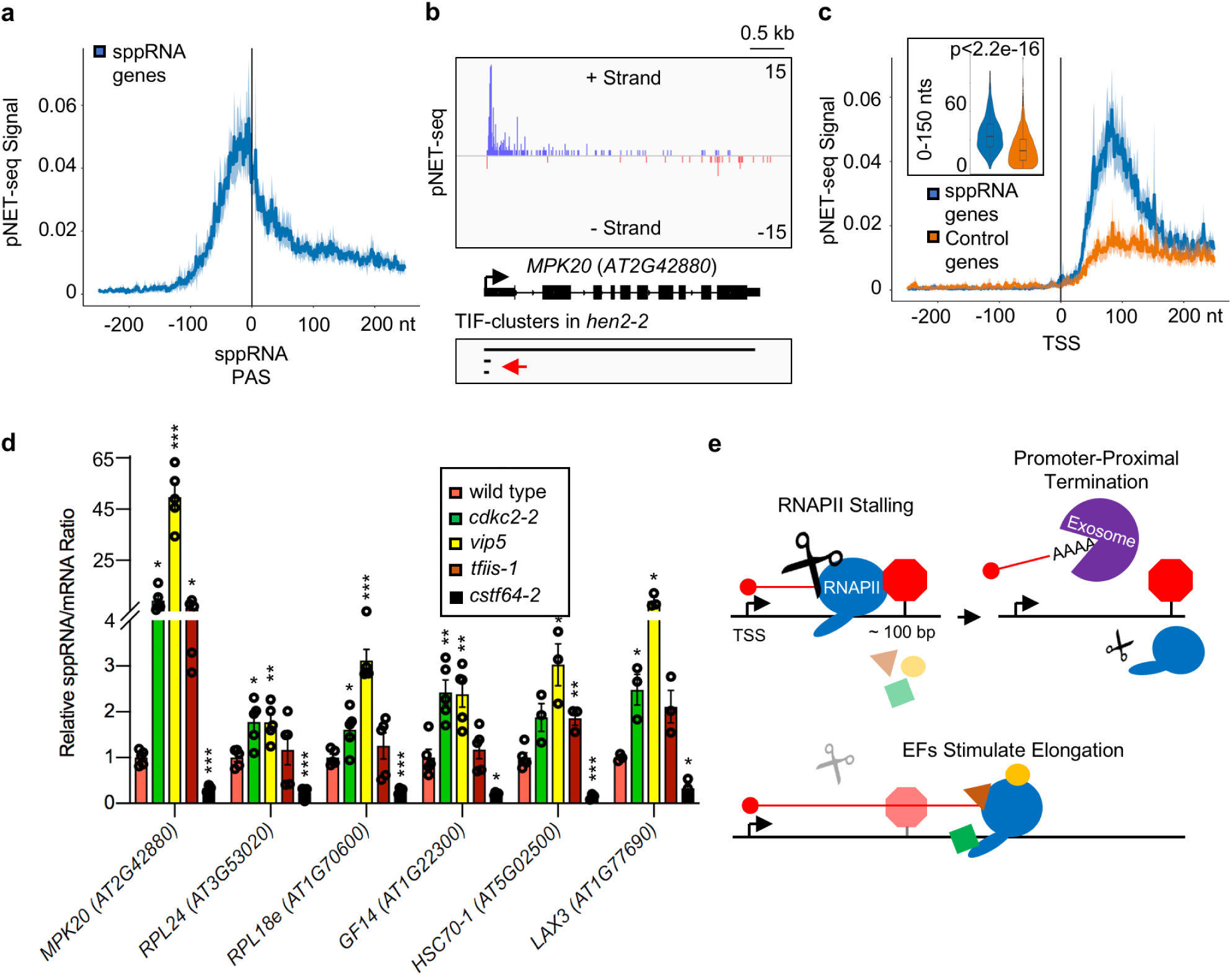
RNAPII stalling coincides with regulated termination near *Arabidopsis* promoters. (a) wild type pNET-seq signal (8WG16)^15^ plotted around sppRNA PAS positions for genes with sppRNAs (n=1155). Shading represents confidence interval of 95%. (b) Genome browser screenshot of wild type pNET-seq reads at the *MPK20* gene. TIF-clusters indicate mRNA and sppRNA positions. (c) pNET-seq signal for genes with sppRNAs (n=1155) and a control set of genes displaying equal gene body transcription (n=1153). See Methods. Inset plot depicts significantly increased average pNET-seq signal within first 150 nt from TSS for genes with sppRNAs (Wilcoxon test: p<2.2e-16). (d) sppRNA/mRNA ratio for six genes with sppRNAs in wild type, *cdkc2-2, vip5, tfiis-1*, and *cstf64-2* by RT-qPCR. Black circles indicate individual data-points. Error bars represent SEM resulting from at least three independent experiments. Single asterisk denotes p<0.05, two asterisks denote p<0.01, while three asterisks denote p<0.001 between mutant and wild type by Student’s t-test. (e) Schematic representation of promoter-proximal RNAPII stalling ∼ 100 nt downstream of TSS and coinciding with nascent transcript cleavage, polyadenylation, and exosome-mediated degradation of sppRNAs. Elongation factors (EF) counter this phenomenon, allowing RNAPII to escape premature termination and synthesize full-length mRNAs.

Elongation factors (EFs) promote gene expression by stimulating transcriptional elongation of RNAPII^16^. For example, Positive Transcription Elongation Factor b (P-TEFb) phosphorylates RNAPII and associated factors to drive elongation^16^. RNAPII peaks at sppRNA genes are particularly sensitive to treatment with the P-TEFb inhibitor Flavopiridol (**Fig. S16**), linking stalled RNAPII to sppRNA formation. To test the connection between RNAPII stalling, P-TEFb, and premature termination, we used an *Arabidopsis* mutant for P-TEFb kinase CYCLIN-DEPENDENT KINASE C2 (CDKC2): *cdkc2-2*^17^. RT-qPCR analyses revealed that sppRNA levels increased in proportion to cognate mRNA levels in *cdkc2-*2 compared to wild type (**Fig. 3d**). Thus, EF activity may be required to bypass a gene expression checkpoint associated with sppRNA formation. We therefore used the sppRNA/mRNA ratio as a quantitative read-out for premature termination in other well-characterized EF mutants. Notably, sppRNA production increased in *vip5* (**Fig. 3d**), a mutant of the *VERNALIZATION INDEPENDENCE 5* (*VIP5*) gene^18^, which encodes a conserved subunit of the Polymerase Associated Factor 1 Complex (PAF1c)^19^. These results suggest *Arabidopsis* PAF1c helps limit premature termination. Transcription factor S-II (TFIIS) stimulates elongation by resolving backtracked RNAPII and enhances RNAPII/PAF1c interactions^20^, yet only some genes tested showed increased sppRNA levels in the *tfiis-1* mutant (**Fig. 3d**)^21^. We next tested if termination failures impact sppRNA levels. We found that disrupting *Arabidopsis* Cleavage Stimulation Factor 64 (CSTF64), a conserved RNA-binding protein involved in pre-mRNA cleavage and polyadenylation, resulted in a significant reduction of sppRNA levels for all genes tested in *cstf64-2* (**Fig. 3d**)^22^. These results suggest *Arabidopsis* EFs promote gene expression by stimulating stalled RNAPII near TSSs to elongate beyond a prominent checkpoint triggering sppRNA cleavage, polyadenylation, and subsequent RNAPII termination (**Fig. 3e**). Collectively, our findings indicate this checkpoint balances bursts of productive full-length isoform production with promoter-proximal termination events to control plant gene expression.

In metazoans, the highest density of RNAPII along genes is observed as “stalled RNAPII” near promoters. These stalled RNAPII complexes may prevent new acts of initiation^23^ and limit full-length isoform production by regulated pause-release mechanisms^24^. Notably, the extent to which stalled RNAPII near metazoan TSSs represent transient transcriptional pausing or dynamic RNAPII turn-over, perhaps through transcriptional termination, is now actively debated^25^. The sppRNA biogenesis mechanism likely differs from shorter RNAs (∼ 20-65 nts) detected near initiation regions that may depend on metazoan pausing factors^24,26-29^. The conspicuous absence of pausing factor homologs in plants^16^ supports transcriptional termination as the outcome of RNAPII stalling near many plant promoters. In sum, our analyses of isoform diversity in plants inform on the debate about peaks of RNAPII occupancy shortly after TSSs, suggesting they often represent RNAPII complexes engaged in premature transcriptional termination.

## Supporting information

TIF-seq cluster statistics

GO term analysis of sppRNA genes

Arabidopsis thaliana lines used in this study

Primers used in this study (Integrated DNA Technologies)

Genomic coordinates of sppRNA

## Acknowledgments

We thank members of the Marquardt and Pelechano laboratories for discussions and technical assistance. We are grateful to T.H. Jensen, S. Buratowski, and P. Brodersen for critically evaluating the manuscript.

## Author contributions

R.A. and S.M. conceived the project. R.A., V.P., and S.M. designed experiments. R.A. performed all experiments. Q.T. performed computational analyses with support from J.W and V.P. B.L., J.W. and V.P. optimized TIF-seq. S.M. supervised the project. R.A. and S.M. wrote the manuscript with input from Q.T.

## Data and materials availability

Data have been deposited to NCBI GEO under accession number GSE129523.

## Supplemental Material

### Materials and Methods

#### Plant growth

All *Arabidopsis thaliana* lines used in this study are listed in **Table S3**. Sterilized *Arabidopsis* seeds were grown on plates containing 1/2 Murashige and Skoog (MS) medium containing 1% sucrose and supplemented with 1% Microagar. Seeds were stratified in the dark at 4°C for 48 hours. Plates were then transferred to climate chambers with a long day photoperiod (16h light/8h dark cycle) at 22°C/18°C and grown for two weeks. Light intensity during control growth conditions was approximately 100 μEm-2s-1. The Columbia accession Col-0 was used as wild type background for all experiments. For cold treatment, seedlings were grown in control conditions for two weeks and subsequently transferred to a cold room (4°C) with light for three hours before harvesting.

#### TIF-seq library construction

Discrete TU boundaries in *Arabidopsis thaliana* were mapped by an optimized version of TIF-seq(Pelechano, Wei et al. 2014). The TIF-seq approach was modified in order to better detect corresponding TSS/PAS pairs in organism more complex than that of budding yeast *Saccharomyces cerevisiae*. Most notably, the make-up and orientation of sequencing adapters was improved to effectively double the return of map-able reads. The detailed protocol will be published elsewhere (Li, Wang *et al.*, In preparation). Here we describe its application to *Arabidopsis thaliana*. Briefly, total *Arabidopsis* RNA was isolated using the RNeasy Plant Mini Kit (Qiagen) and treated with Turbo DNase (Thermo Fisher Scientific) according to manufacturers’ instructions. DNase-treated RNA was recovered by acid-phenol extraction and ethanol precipitation. RNA integrity was assessed using the 2100 Bioanalyzer RNA 6000 Nano assay (Agilent). 10 micrograms of DNase-treated total RNA was treated with CIP (NEB) according to manufacturer’s instructions in order to remove all non-capped RNA species in the sample. RNA was recovered by acid-phenol extraction and ethanol precipitation. Next, 5’ caps were removed using Cap-Clip (CellScript). RNA was quickly recovered by acid-phenol extraction and ethanol precipitation. The single-stranded rP5_RND adapter (see **Table S4** for oligonucleotide sequence) was ligated to the 5’-end of previously capped species using T4 RNA ligase 1 (NEB). RNA was recovered by AMPure purification using Agencourt RNAClean XP beads (Beckman Coulter) following manufacturer’s instructions. Before proceeding, RNA integrity was assessed using the 2100 Bioanalyzer RNA 6000 Nano assay (Agilent) following manufacturer’s instructions. In order to generate full length cDNAs up to and greater than 10 kb long, SuperScript IV (Invitrogen) and an oligo(dT)-containing barcoded adapter (see **Table S4** for TIF2-RTX oligonucleotide sequences) was used with the following PCR steps: 10 mins at 42°C, 30 mins at 50°C, 30 mins at 55°C, and 10 mins at 80°C. cDNA samples were treated with RNase H, provided with SuperScript IV First-Strand Synthesis System (Invitrogen), and recovered using AMPure XP beads (Beckman Coulter). Second-strand synthesis of full length cDNA was amplified with Terra PCR Direct Polymerase Mix (Takara) the BioNotI-P5-PET oligo (see **Table S4** for oligonucleotide sequence) and the following PCR steps: 2 mins at 98°C, 10 cycles (20 secs at 98°C, 15 secs at 60°C, 5 mins at 68°C + 10 secs/cycle), 5 mins at 72C. Barcoded full length double-stranded cDNA libraries were recovered using AMPure XP beads (Beckman Coulter) and assessed for quality using the 2100 Bioanalyzer High Sensitivity DNA kit (Agilent). cDNA libraries were quantified with the Qubit dsDNA HS Assay Kit (Thermo Fisher Scientific) according to manufacturer’s instructions. Equal quantities of cDNA libraries (up to 600 ng total) were pooled and digested with NotI (NEB) for 30 mins at 37°C. The enzyme was inactivated at 65°C. Digested cDNA was recovered using AMPure XP beads (Beckman Coulter). Digested linear cDNA pools were diluted to less than 1 ng/ul and circularized using T4 DNA ligase (NEB) at 16°C overnight. PlasmidSafe™ DNase (Cambio) was added for 30 min at 37°C to remove linear DNA molecules from the sample and then inactivated at 70°C for 30 min. Circular cDNA molecules were recovered by phenol-chloroform extraction followed by isopropanol precipitation. Circular cDNA was sheared by sonication at 4°C in a Bioruptor (Diagenode) (3 cycles, 30s ON, 90s OFF). Sheared DNA fragments were purified with AMPure XP beads (Beckmann Coulter). Biotinylated fragments containing flanking TSS/PAS pairs were captured with M-280 Streptavidin Dynabeads (Thermo Fisher Scientific), end repaired with End Repair Enzyme mix (NEB), A-tailed with Klenow fragment exo-(NEB), and ligated to a common Illumina-compatible forked adapter using T4 DNA ligase (NEB) according to manufacturers’ instructions. Libraries were amplified using the Phusion High-Fidelity PCR mix (NEB) with the following PCR steps: 98°C for 1 min, 15 cycles (20 secs at 98°C, 30 secs at 65°C, 30 secs at 72°C), 5 mins at 72C. Libraries were size selected using AMPure XP beads (Beckman Coulter) and sequenced with the following flowcell: FC-404-2002 NextSeq 500/550 High Output v2 kit (150 cycles) (Illumina). The sequencing run was carried out using a mix 1:1 of custom oligos SeqR1+15T and SeqR2 for sequence reads 1 and 2, and of custom oligos SeqINDX and TIF3-RvRT for the index reads 1 and 2 (see **Table S4** for oligonucleotide sequences). Cluster orientation was determined based on the information contained in the index reads. All TIF-seq experiments were performed as independent duplicates.

#### TSS-seq library construction

TSSs were mapped genome-wide in *Arabidopsis* using 5’-CAP-sequencing(Pelechano, Wei et al. 2016), with minor changes previously described(Kindgren, Ard et al. 2018, Nielsen, Ard et al. 2019). Note, we define this adapted method Transcription Start Site sequencing (TSS-seq). Here, we extend our previous TSS-seq analyses to two-week old *hen2-2* seedlings before and after cold-treatment (3hours at 4°C) in order to correct for TIF-TSS bias (see Computational methods below). All TSS-seq experiments were performed as independent duplicates.

#### RT-qPCR

RNA was isolated from two-week old *Arabidopsis* seedlings using Plant RNeasy Mini-Kits (Qiagen) and treated with Turbo DNase-treated (Ambion) as per manufacturers’ instructions. For selective quantitative analysis of mRNAs and sppRNAs, first strand complementary DNA (cDNA) synthesis was performed on RNA using an oligo(dT)-containing primer (see **Table S4** for oligonucleotide sequence) and SuperScript IV First-Strand Synthesis System (Invitrogen) following manufacturer’s instructions. cDNA samples were treated with RNase H (Invitrogen) for 20 mins at 37°C before enzyme inactivation at 80°C for 20 mins. Negative controls lacking the reverse transcriptase enzyme (-RT) were performed alongside all RT–qPCR experiments. Quantitative analysis of mRNAs (F1/R1 primer pairs) or sppRNAs (F1/R2 primer pairs) were performed by qPCR using the CFX384 Touch Real-Time PCR Detection System (Biorad) and GoTaq qPCR Master Mix (Promega). See **Table S4** for oligonucleotide sequences used. Short annealing and extension times were employed to specifically amplify and distinguish sppRNAs from their cognate mRNAs. qPCR cycles were as follows: 5 mins at 98°C, 40 cycles (10 secs at 98°C, 5 secs at 55°C, 5 secs at 65°C). Data was normalized to the mRNA levels of an internal reference gene lacking detectable sppRNAs (*ACT2*). To compare relative sppRNA/mRNA changes, *sppRNA*/*ACT2* levels were first normalized to the expression of cognate *mRNA*/*ACT2* levels. Next, the sppRNA/mRNA ratio calculated in different mutants was expressed relative to the sppRNA/mRNA ratio calculated in wild type. Error bars reflect SEM resulting from at least three independent replicates. See **Table S5** for values used to construct plots.

#### Northern blotting

Northern analysis was performed as previously described(Nielsen, Ard et al. 2019) with minor modifications. Briefly, 5 micrograms of total RNA was separated by electrophoresis on agarose-formaldehyde-MOPS gels and transferred to a nylon transfer membrane by capillary blotting in 6x SSC overnight. RNA was crosslinked to the nylon membrane by UV irradiation using a Stratalinker UV Crosslinker (Stratagene). Membranes were prehybridized in Church Buffer (0.5 M Na_2_HPO_4_ pH 7.2, 1% BSA, 1 mM EDTA, 7% SDS) for 1 hour at 68°C. Next, membranes were probed overnight at 68°C with single stranded DNA probes generated by PCR-based amplification and incorporation of radioactive α-32P-dTTP (PerkinElmer). Membranes were washed twice in pre-warmed buffer containing 2X SSC and 0.1% SDS for 10 mins (68°C) before exposure to Storage Phosphor Screens (Kodak). A Typhoon phosphoimager (GE Healthcare Life Sciences) was used for analysis. For loading controls, membranes were stripped three times with a near boiling buffer containing 0.1X SSC and 0.1% SDS before prehybridization in Church Buffer as above and northern blotting against the mRNA of a reference gene lacking detectable sppRNAs (*UBI*). See **Table S4** for oligonucleotide sequences used to prepare radioactive DNA probe templates.

#### Bioinformatics

Computational analysis of TSS-seq data analysis was performed as previously described(Nielsen, Ard et al. 2019). For TIF-seq, paired-end sequencing reads were demultiplexed and concatenated according to the indexes adjacent to 5’- or 3’-ends of transcript molecules. Reads were then subjected to adapter- and UMI-trimming with respectively cutadapt v1.18 (AGGTGACCGG, AGATCGGAAG) and UMI-Tools v0.5.5. Extra stretches of As near the polyadenylation sites were removed from reads using cutadapt v1.18. Next, reads containing 10 nt long stretches of As and Ts with one mismatch were removed using AfterQC v0.9.7. The adapter-, UMI- and polyX-trimmed reads were aligned to TAIR10 genome assembly using STAR v2.6.1c (--alignEndsType EndToEnd --alignMatesGapMax 15000). The output sorted BAM files were filtered for uniquely aligned reads using SAMtools. Finally, the filtered BAM files were deduplicated using UMI-Tools dedup.py. The two replicate alignments of each data-set were compared using deepTools multiBamsummary, calculating a coverage matrix between replicates (--binSize 10). The pearson correlation for each matrix was plotted with the deepTools plotCorrelation (--corMethod pearson --log1p and –removeOutliers) in **Fig. S3b, Fig. S4a/c, Fig. S5a, Fig. S11a** and **Fig. S12a**, showing reproducibility between replicates. Based on the reproducibility, the two replicates of TIF-Seq were merged before clustering with SAMtools.

Each pair of reads aligned to the genome was clustered with the custom script “TIF-Seq_isoform_clustering_v6.py” as follows: “Pre-clusters” were called and merged read pairs overlapping exactly at both ends. Next, Pre-clusters were merged if both start and end positions overlapped another existing “Pre-cluster” by <20 nucleotides. Subsisting merged pre-clusters were kept as final clusters only if their pairs of reads count was equal or higher than 3. The remainder were discarded. To further remove clusters produced by accidental 3’-end mis-priming, extended clusters overlapping genomic polyA regions (from 6 to 9 As) were called with bowtie aligner and discarded. Due to artefactual bias of TIF-seq called TSSs towards genic 3’-ends, as previously described(Pelechano, Wei et al. 2014), the 5’ reads of TIF-Seq were used to call TIF-TSSs for each dataset using the CAGEfightR package v1.0.0 package that is available from Bioconductor (https://bioconductor.org/packages/release/bioc/html/CAGEfightR.html) and filtered out if they did not overlap a previously defined TSS called from published TSS-Seq data(Nielsen, Ard et al. 2019). The density of TSSs called with CAGEfightR for TSS-Seq and TIF-Seq in wild type, including the TIF-TSSs bias towards small clusters overlapping the end of genes, is shown in **Fig. S3a**. The overall numbers of paired reads along with discarded and final TIF-clusters called per genotype can be found in **Table S1**.

For all further genome-wide clusters analysis, measurements considered only non-overlapping genes to avoid false positive calculations while accepting to lose some information. In **Fig. 1c, Fig. 2a, Fig. S11d** and **Fig. S12e** we calculated the exact distance between each cluster boundary (from the final TIF-clusters) to their corresponding overlapping gene boundaries extracted from TAIR10 coding genes annotation available through the TxDb.Athaliana.BioMart.plantsmart28 R package. then plotted using the “comparisonplot()” function from LSD R package (cf script “histscatter_all.R”). In the case of **Fig. 2c** and **Fig. S12g**, the same calculation was performed in *hen2-2* but only on clusters with widths <200 nucleotides (cf script “histscatter_small.R”). Finally, the same distance calculation was performed for **Fig. 1f** on a subset of TIF-Seq clusters in FACT mutants *ssrp1-2* and *spt16-1* of which starts overlapped with known *fact*-specific TSSs(Nielsen, Ard et al. 2019) (cf script “histscatter_FACT.R”). To evaluate the distribution of clusters sizes between samples, all clusters <6000 nucleotides in width were plotted as a violin plot between wild type and *hen2-2* in **Fig. 2b**. To establish the positional distribution of isoforms across genes, the clusters boundaries were overlapped with the corresponding genes boundaries and segregated into different classes (TIF Cluster Categories) with specific comparison to annotated genes using the countOverlaps() function from R package “Genomic Ranges” (cf script “sepclusters_orf_simplified.R” and “sepclusters_overlapping_orf.R”). Each number for TIF cluster categories were compared to the ensemble of all genes overlapping TU clusters. These analyses are represented as piecharts in **Fig. 1d, Fig. S4e-f, Fig. S5c, Fig. S11c**, and **Fig. S12c**. The log2 Fold Change of the proportion of each category across genotype in wild type and *hen2-2* was calculated in **Fig. S5d** and between cold-treated wild type and *hen2-2* in **Fig. S12f.** The number of clusters overlapping the whole ORF was measured for each gene in each dataset and plotted as histograms in **Fig. S3c, Fig. S4b/d** and **Fig. S5b.**

To extract small promoter-proximal clusters, all clusters <350 nucleotides were mapped to annotated TSSs extended by 40 nucleotides upstream and downstream and to overlap no termination sites to account for TSS variation. The corresponding set of overlapping clusters were then called short promoter-proximal RNAs (sppRNAs; see **Table S6** for full list of genomic coordinates). sppRNA size distribution was plotted in **Fig. S5e** (cf script “sepclusters_forsmallclusters.R”).

All further analysis requiring expression controls were calculated using transcription profiles of published unphosphorylated RNAPII (8WG16) pNET-Seq wild type datasets(Zhu, Liu et al. 2018). pNET-Seq signal was averaged over all genes shrunk by 200 nucleotides downstream of the TSS (+200 from TSS) and 200 nucleotides upstream of the PAS (−200 from TSS) to reduce biases from recently reported peaks in nascent transcription signals near TSS-proximal or PAS-proximal positions(Zhu, Liu et al. 2018). The percentage of cold-induced genes (calculated from wild type TSS-seq data) that only have sppRNAs at 22°C was compared against a control set of genes without detectable sppRNAs in **Fig. S13B**. In **Fig. S13c**, all sppRNA-containing genes common between wild type and *hen2-2* datasets before and after cold treatment were plotted as a Venn Diragram (cf script “ViennDiagram.R”).

The distribution of nascent transcriptional activity over sppRNA genes was compared to all other genes with no sppRNAs in wild type in **Fig. S6**. A set of genes with no sppRNAs detected in any dataset and with equal distribution of nascent gene body transcription (pNET-seq) was calculated using the script “subset_equaldistribution_v4.R” and used as control set of genes. TSS positions were used as anchor points in the metagene plot found in **Fig. 3c** and **Fig. S16a-c** as described(Nielsen, Ard et al. 2019). In **Fig. 3a**, just the last position of sppRNAs (i.e. the PAS) were used as an anchor for the metagene plot. The proportion of genes with sppRNAs in relation to the 5’ pausing index value in **Fig. S15** was plotted for 12,854 non-overlapping genes longer than 1000 nt and belonging to the 75% most expressed genes. The 5’ pausing index value was calculated as previously described(Zhu, Liu et al. 2018) by dividing the promoter TPM coverage from pNET-seq (150 nt upstream and downstream arround the TSS) to the gene body (shrunk gene region by 300 nt downstream of the TSS and 300 nt upstream of the PAS).

Finally, all genome browser screenshots were made using Integrative Genomics Viewer (IGV_2.3.93) with the *Arabidopsis thaliana* (TAIR 10) annotation. In particular, TIF-seq reads (BAM alignment files, before clustering) were visualized with the following settings: collapsed, view as pairs, colour alignments by read strand.

#### Data availability

TIF- and TSS-seq data is available at NCBI GEO database with accession code GSE129523: https://www.ncbi.nlm.nih.gov/geo/query/acc.cgi?acc=GSE129523. TIF- and TSS-seq analysis used computational methods. Scripts for these analyses can be found at github: https://github.com/qthom/planTIF. In github, bed files representing every read pair after filtration but not clustering is available to be uploaded into a genome browser. Source data underlying figures can be found in **Table S5**, while genomic coordinates for sppRNA TIF-clusters are listed in **Table S6**. All other data supporting the findings of this study are available from the corresponding author upon request.

## Supplementary Figures and Legends

**Figure S1.**
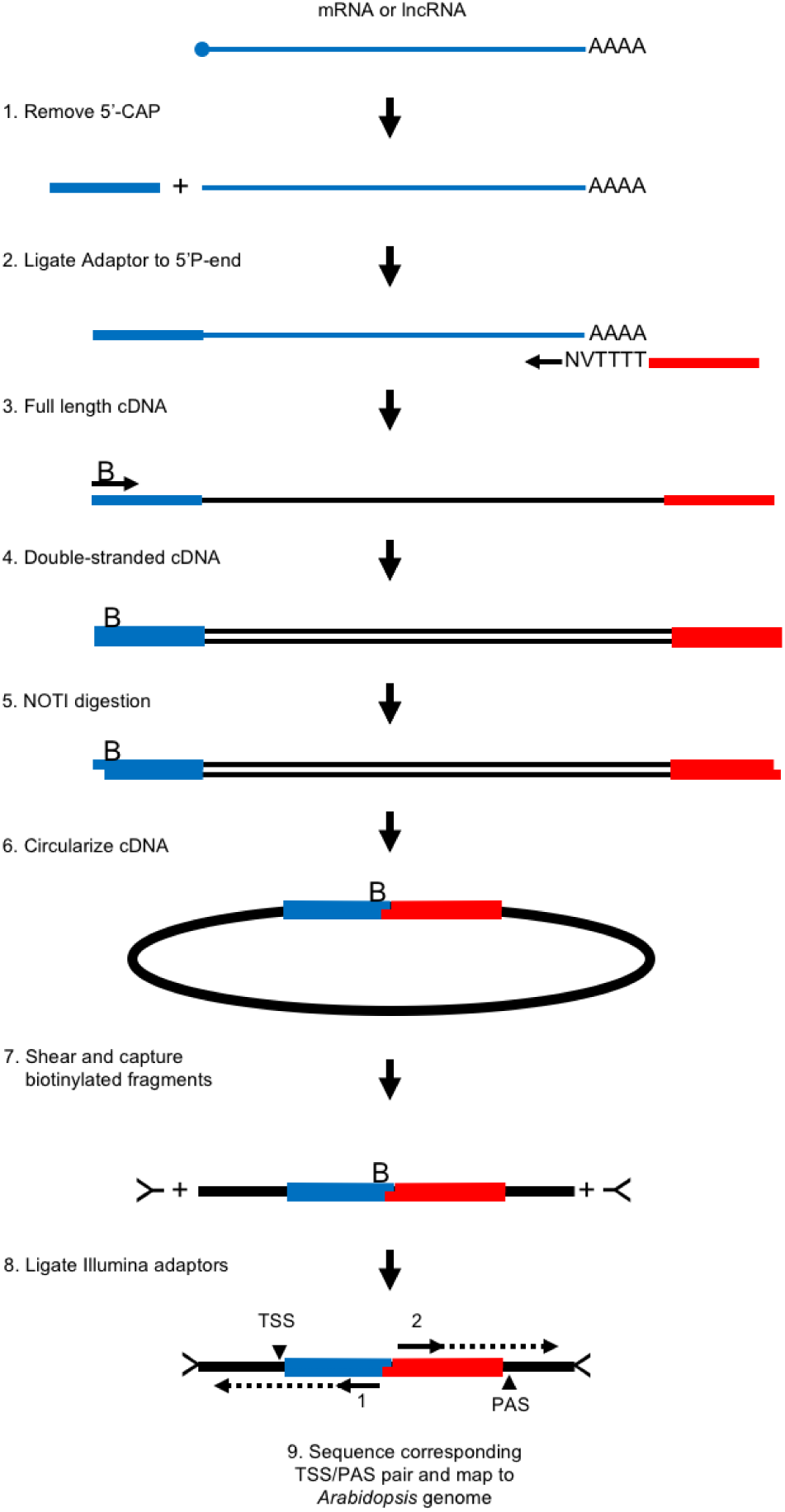
Improved TIF-seq method workflow. Schematic representation of TIF-seq workflow (see Methods). A notable change to the original method include custom Illumina-compatible 5’ and 3’ adaptors for sequencing out of the insert. For other improvements to the protocol, see Methods. The letter “B” indicates a biotin-group added during double-stranded cDNA synthesis step for streptavidin capture. First and second reads identify TIF-TSS and TIF-PAS, respectively.

**Figure S2.**
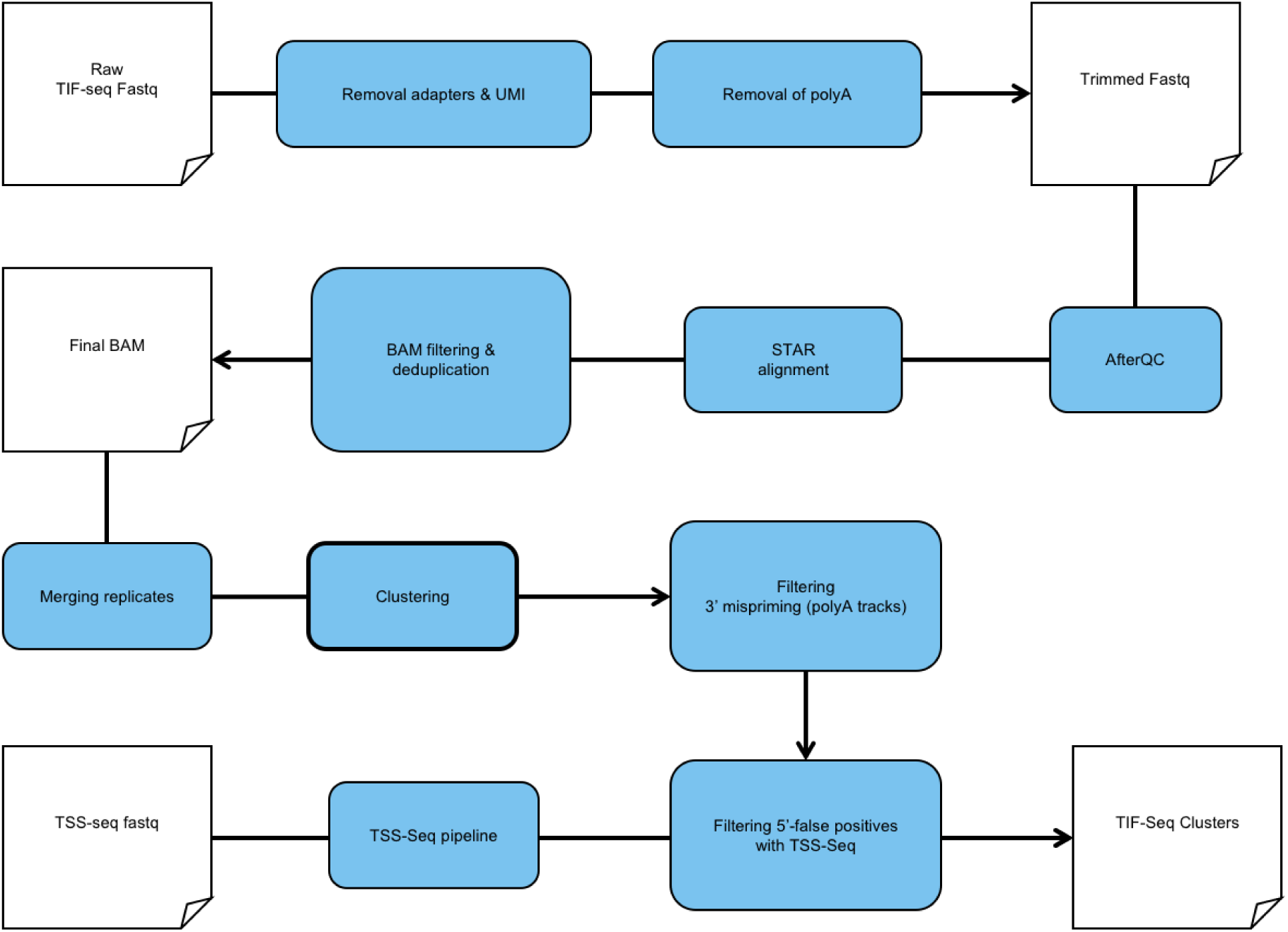
Computational workflow for TIF-seq data. Schematic representation of workflow for handing TIF-seq data and calling TIF-clusters (see Methods). The pipeline for handling TSS-seq data has been previously described(Kindgren, Ard et al. 2018, Nielsen, Ard et al. 2019).

**Figure S3.**
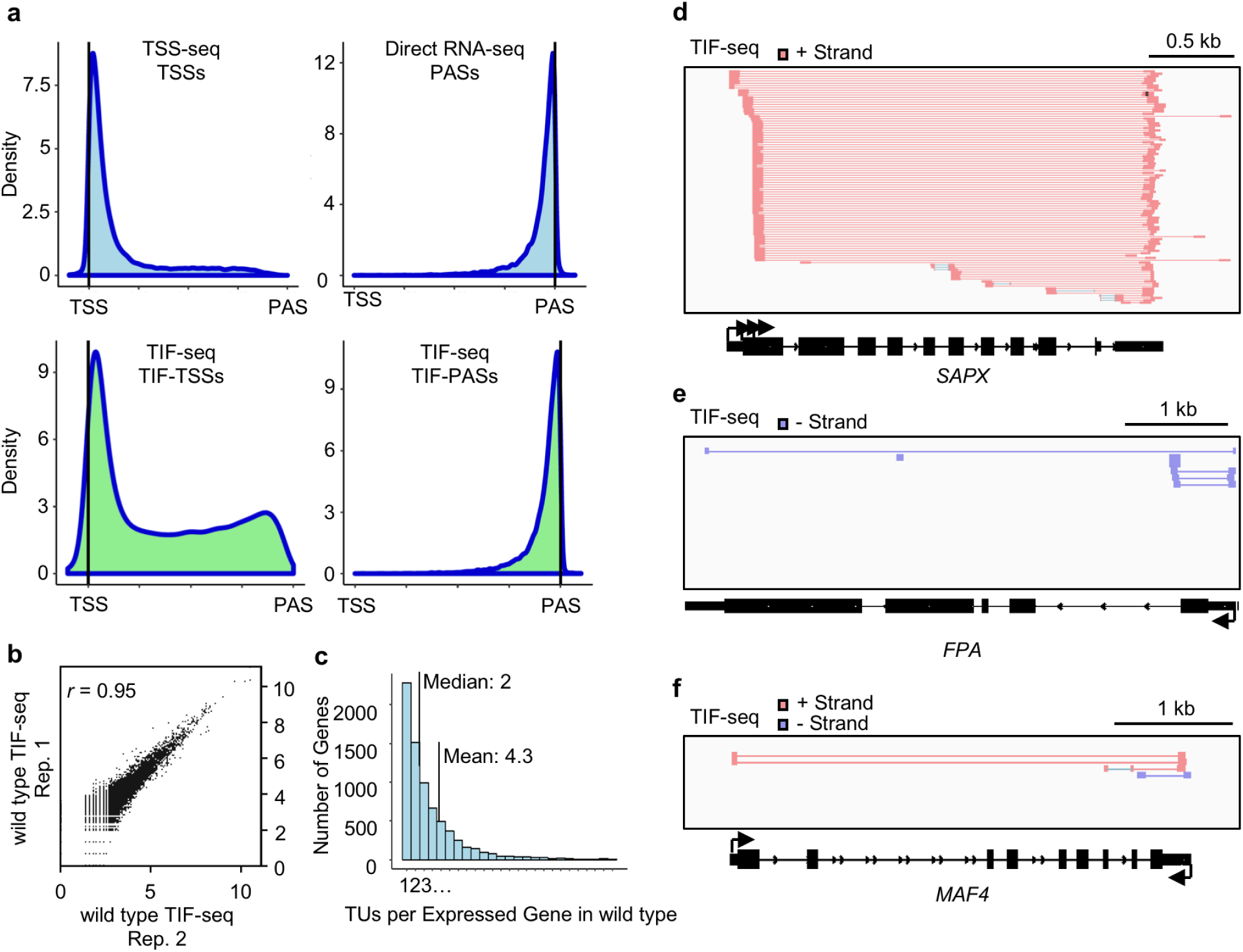
TIF-seq control analyses for wild type validate characterized alternative gene isoforms. (a) Density plots for TSS positions from TSS-seq(Nielsen, Ard et al. 2019) and TIF-seq (TIF-TSSs) and PAS positions from Direct RNA-seq(Schurch, Cole et al. 2014) and TIF-seq (TIF-PASs). Bias of TIF-Seq to detect short molecules, and thus enriched on internal cryptic transcpts (TSS biased towards the 3’ end), has been previously reported(Pelechano, Wei et al. 2014). This can be corrected by excluding TIF-clusters without a TIF-TSS validated by a second method, such as TSS-seq. 75.8% of TIF-clusters in wild type were validated by TSS-seq, while the remaining 14.2% (mostly enriched towards genic 3’-ends) were discarded. See **Table S1** and Methods for more details. See Figure 1C for the distribution of TIF-TSSs following this data processing step. (b) Scatterplot of log_10_(coverage)/5 nucleotide bins for TIF-reads from two independent replicates in wild type (see Methods). (c) Histogram of number of TIF-clusters reveals a mean of 4.3 TUs and a median of 2 TUs per expressed gene in wild type. Genome browser screenshots of TIF-reads for previously characterized alternative gene isoforms stemming from (d) alternative TSSs at the *STROMAL ASCORBATE PEROXIDASE* (*SAPX)* gene(Chew, Whelan et al. 2003), (e) alternative PAS at the *FLOWERING PROTEIN A* (*FPA*) gene(Hornyik, Terzi et al. 2010), and (f) a recently characterized antisense transcript that regulates the *MADS AFFECTING FLOWERING 4* (*MAF4*) gene(Zhao, Li et al. 2018).

**Figure S4.**
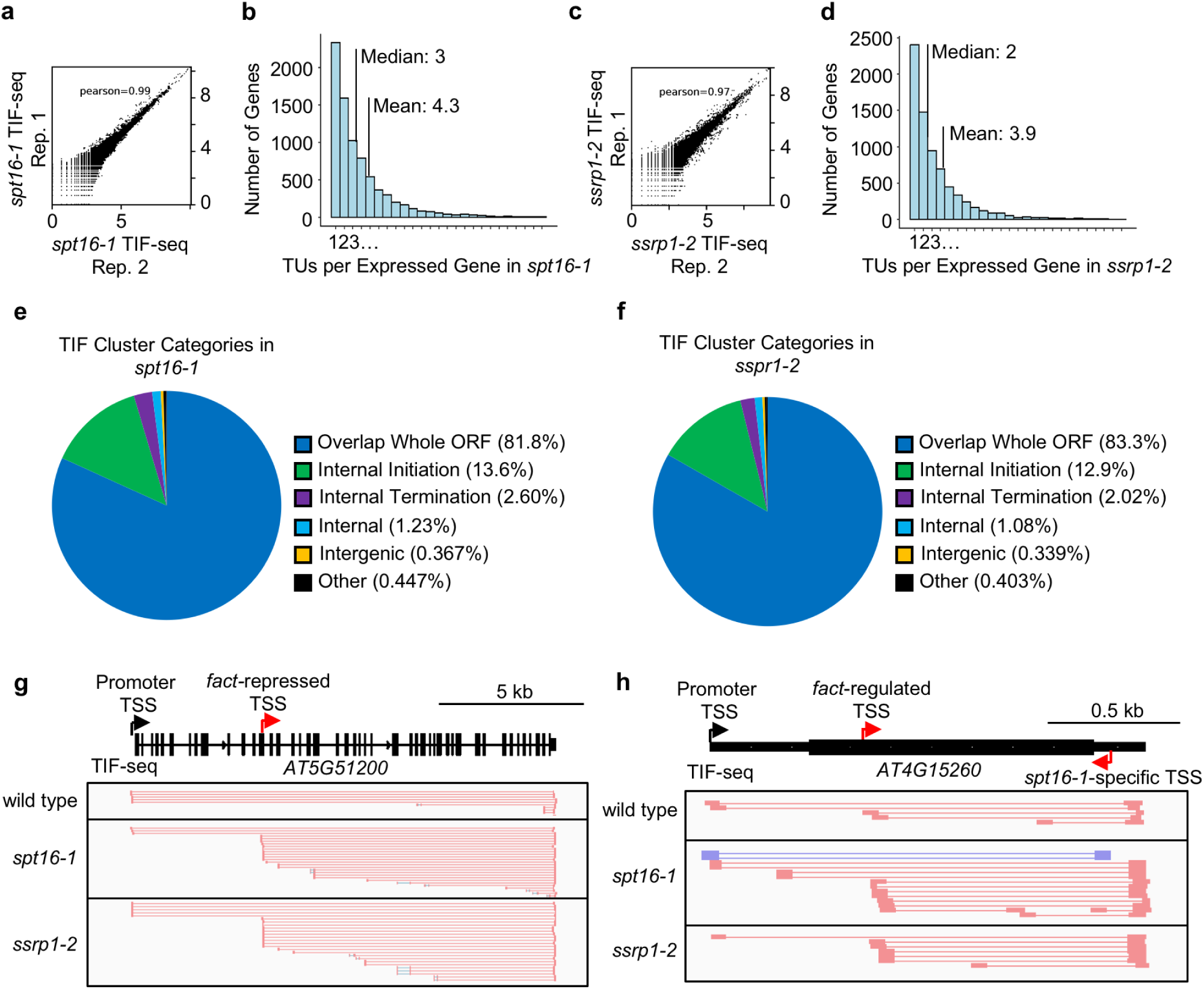
TIF-seq analyses in *fact* mutants reveal co-transcriptional regulation of alternative gene isoforms. FACT consists of two subunits, Suppressor of Ty 16 (SPT16) and Structure-specific recognition protein 1 (SSRP1). (a) Scatterplot of log_10_(coverage)/5 nt bins for TIF-reads from two independent replicates in FACT mutant *spt16-1* (see Methods). (b) Histogram of number of TIF-clusters per expressed gene in *spt16-1* (mean: 4.3; median: 3). (c) Scatterplot of log_10_(coverage)/5 nt bins for TIF-reads from two independent replicates in FACT mutant *ssrp1-2* (see Methods). (d) Histogram of number of TIF-clusters per expressed gene in *ssrp1-2* (mean: 3.9; median: 2). Pie-charts displaying the proportion of TIF Cluster Categories detected in *fact* mutants: (e) *spt16-1* and (f) *ssrp1-2*. Genome browser screenshot of TIF-reads in wild type and *fact* mutants (*spt16-1* and *ssrp1-2*) depicting activation of FACT-repressed intragenic TSSs in *fact* mutants, including at the (g) *AT5G51200* gene, which possesses a recently identified alternative FACT-repressed exonic TSS, and at the (h) *AT4G15260* gene, which possesses a light and FACT-regulated exonic TSS and an *spt16-1*-specific antisense TSS in the 3’UTR(Nielsen, Ard et al. 2019).

**Figure S5.**
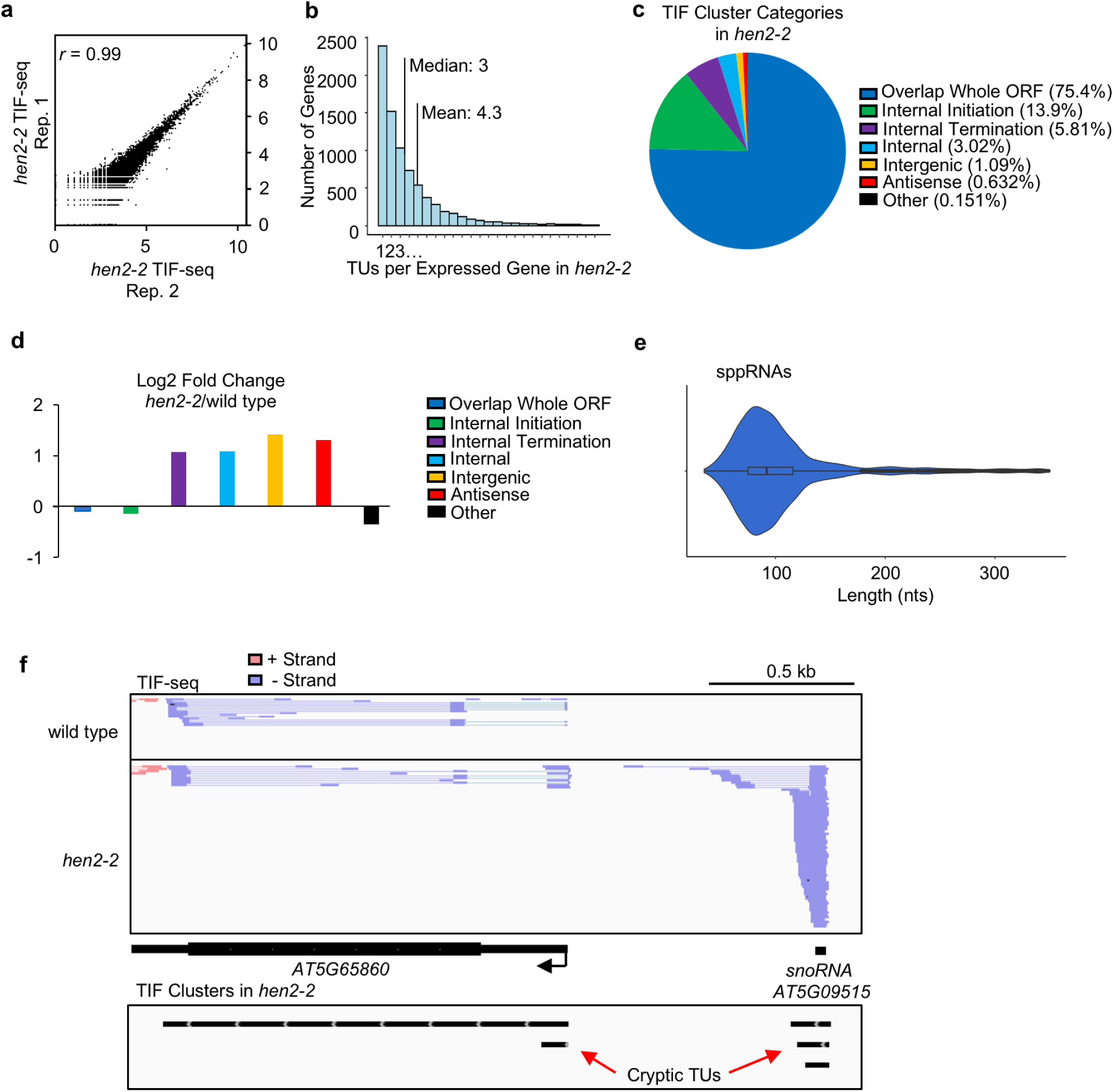
TIF-seq analyses in *hen2-2* reveal cryptic TU boundaries genome-wide in *Arabidopsis*. (a) Scatterplot of log_10_(coverage)/5 nt bins for TIF-reads from two independent replicates in *hen2-2* (see Methods). (b) Histogram of number of TIF-clusters per expressed gene in *hen2-2* (mean: 4.3; median: 3). (c) Pie-chart displaying the proportion of TIF Cluster Categories detected in *hen2-2*. (d) log2 fold change of TIF Cluster category proportions in *hen2-2* compared to wild type. (e) Distribution of short TIF-clusters sizes (<350 nts). sppRNAs have a median length of 93 nt. Cryptic TUs accumulate in *hen2-2*, including snoRNA precursors(Lange, Zuber et al. 2014). (f) snoRNA precursors are visible upstream of *AT5G65860* in *hen2-2*.

**Figure S6.**
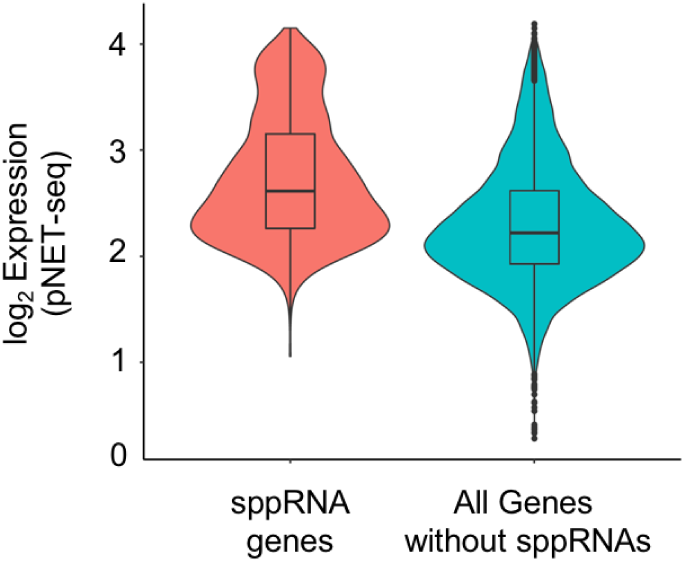
sppRNAs are derived from genes with relatively high gene body transcription. The distribution of nascent gene body RNAPII transcriptional activity for genes with sppRNAs compared to all other genes from pNET-seq (8WG16) in wild type *Arabidopsis*(Zhu, Liu et al. 2018). For both classes tested: n= 1639. Gene body is defined as +200 nts from annotated TSS and −200 nts from annotated PAS. See Methods.

**Figure S7.**
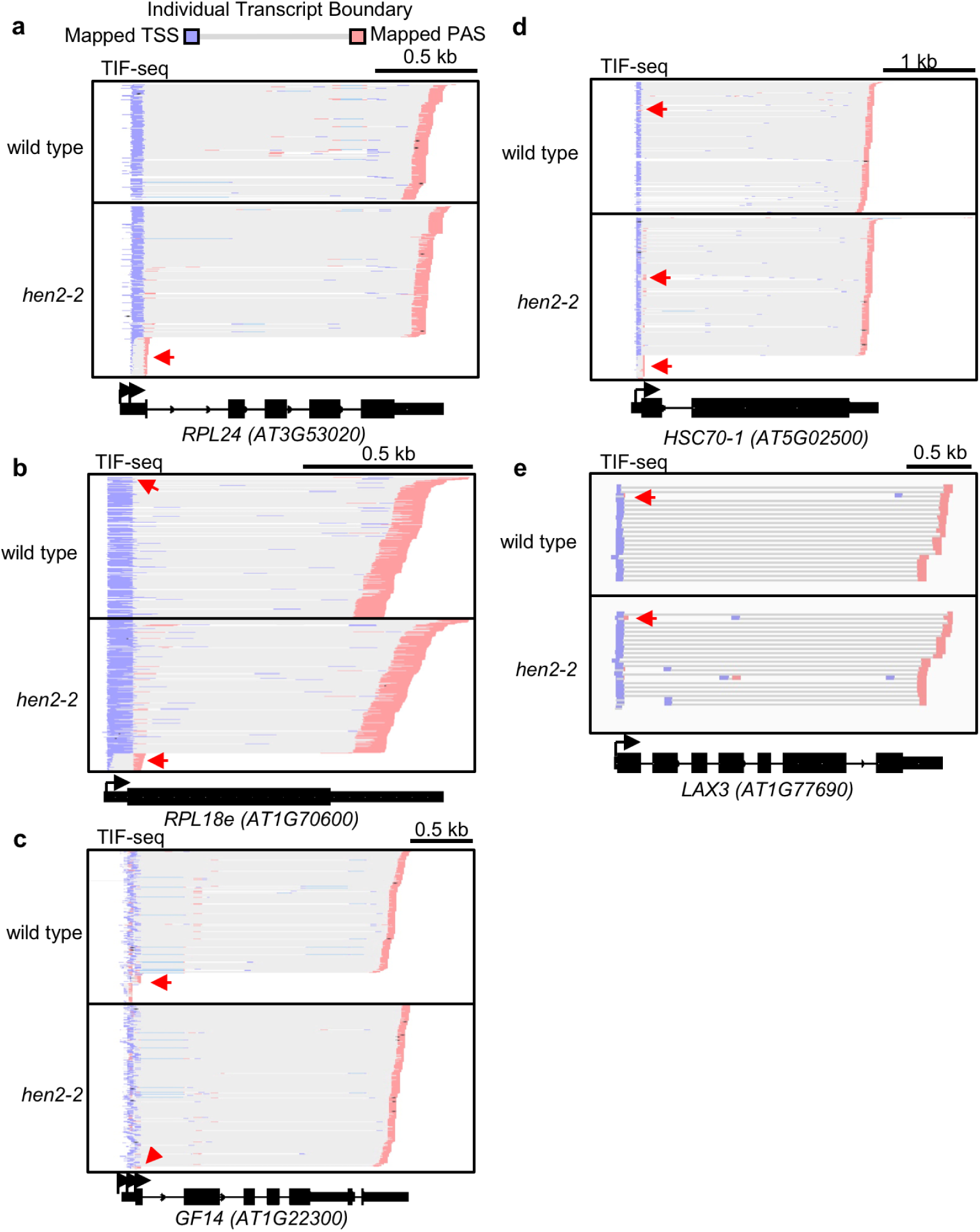
Genome browser screenshots of sppRNAs at genes with varying stead-state mRNA levels. Genome browser screenshots of TIF-reads for additional genes with detected sppRNA TIF-clusters used here to study premature-termination in *Arabidopsis*: (a) *RIBOSOMAL PROTEIN L24* (*RPL24; AT3G53020*), (b) *RIBOSOMAL PROTEIN L18e/L15* (*RPL18e; AT1G70600*), (c) *GENERAL REGULATORY FACTOR 14 EPSILON* (*GF14; AT1G22300*), (d) *HEAT SHOCK COGNATE PROTEIN 70-1* (*HSC70-1; AT5G02500*), and (e) *LIKE AUX1 3* (*LAX3*; *AT1G77690*). Red arrows indicate termination within initiation regions.

**Figure S8.**
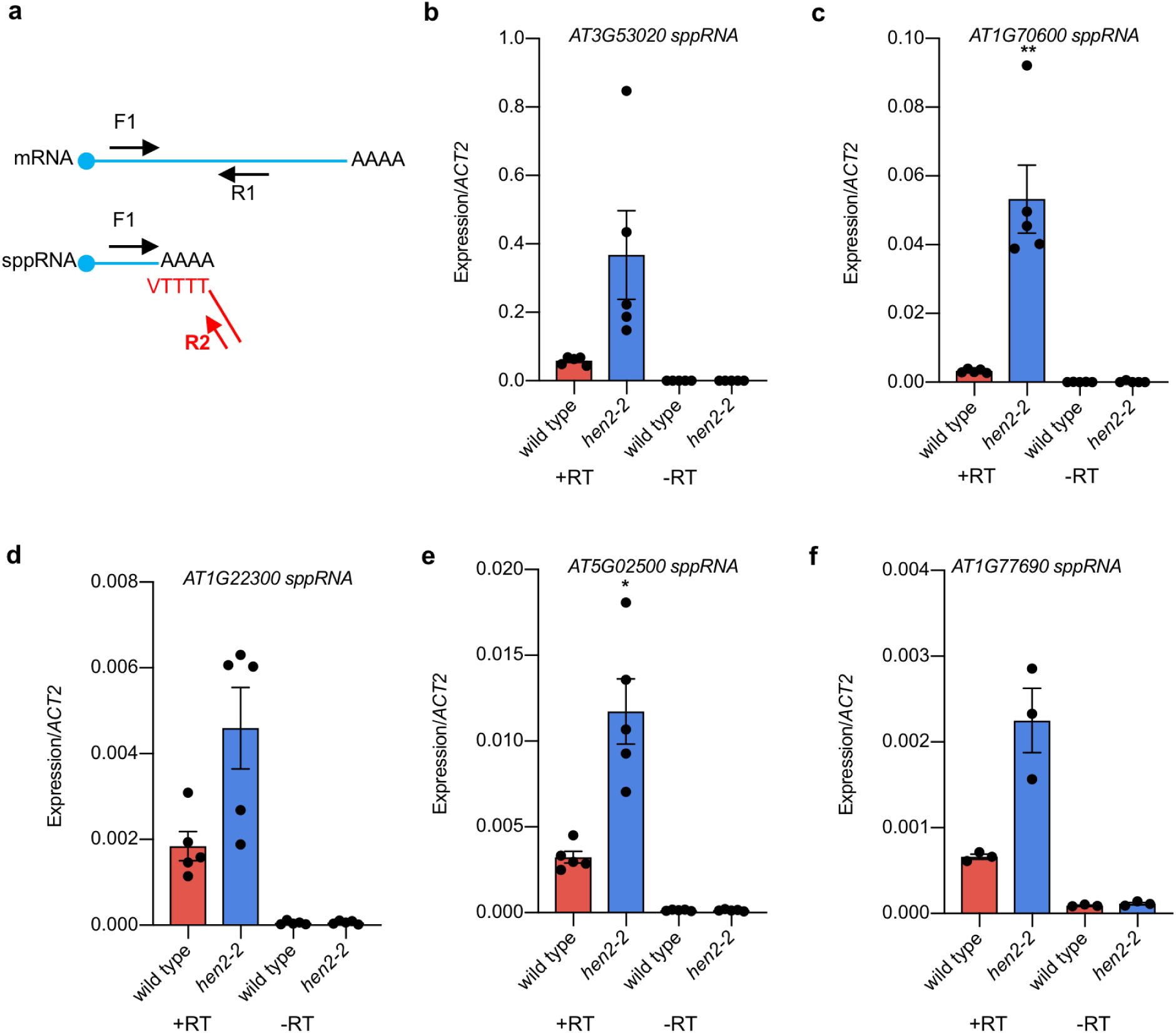
Quantifying sppRNA levels in wild type and *hen2-2*. (a) Schematic representation of RT-qPCR-based strategy to selectively amplify sppRNAs using short annealing and extension times (see Methods). RT-qPCR analysis of sppRNA levels (using F1/R2 primer pairs) detected in wild type and *hen2-2* for (b) *RPL24 (AT3G53020)*, (c) *RPL18e (AT1G70600)*, (d) *GF14 (AT1G22300)*, (e) *HSC70-1 (AT5G02500)*, and (f) *LAX3* (*AT1G77690)*. Error bars represent SEM resulting from at least three independent experiments. Reactions with reverse transcriptase enzyme (+RT) and without (-RT, negative control). Black dots indicate individual data points from independent experiments. For statistical test, a single asterisk denotes p<0.05, while two asterisks denotes p<0.01, between wild type and *hen2-2* by Student’s t-test.

**Figure S9.**
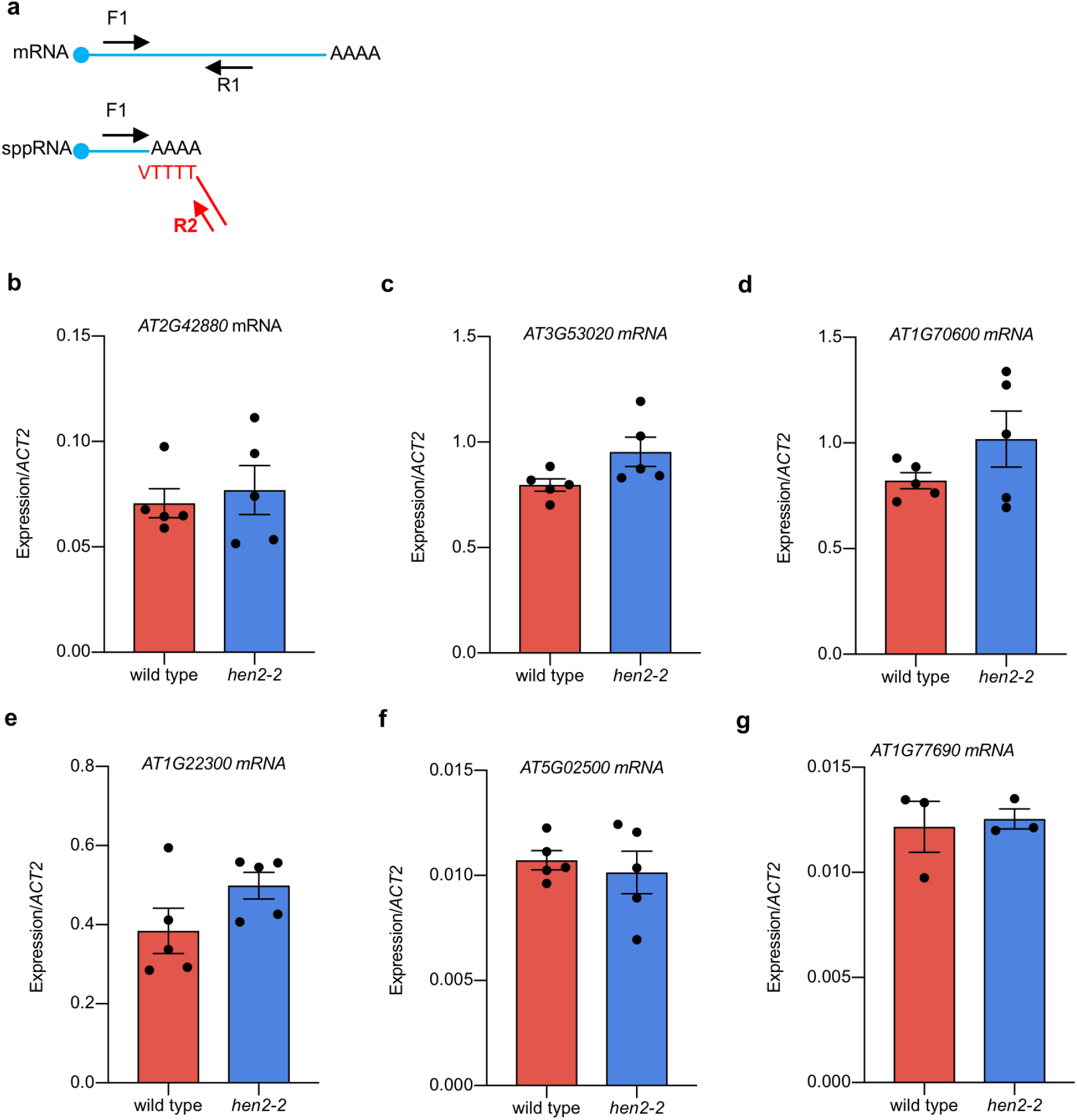
Stabilizing sppRNAs in *hen2-2* does not significantly change cognate mRNA expression. (a) Schematic representation of RT-qPCR-based strategy to selectively amplify sppRNAs and mRNAs using very short annealing and extension times (see Methods). RT-qPCR analysis of mRNA levels (using F1/R1 primer pairs) detected in wild type and *hen2-2* for (b) *MPK20 (AT2G42880)*, (c) *RPL24 (AT3G53020)*, (d) *RPL18e (AT1G70600)*, (e) *GF14 (AT1G22300)*, (f) *HSC70-1 (AT5G02500)*, and (g) *LAX3* (*AT1G77690)*. Black dots indicate individual data points. Error bars represent SEM resulting from at least three independent experiments.

**Figure S10.**
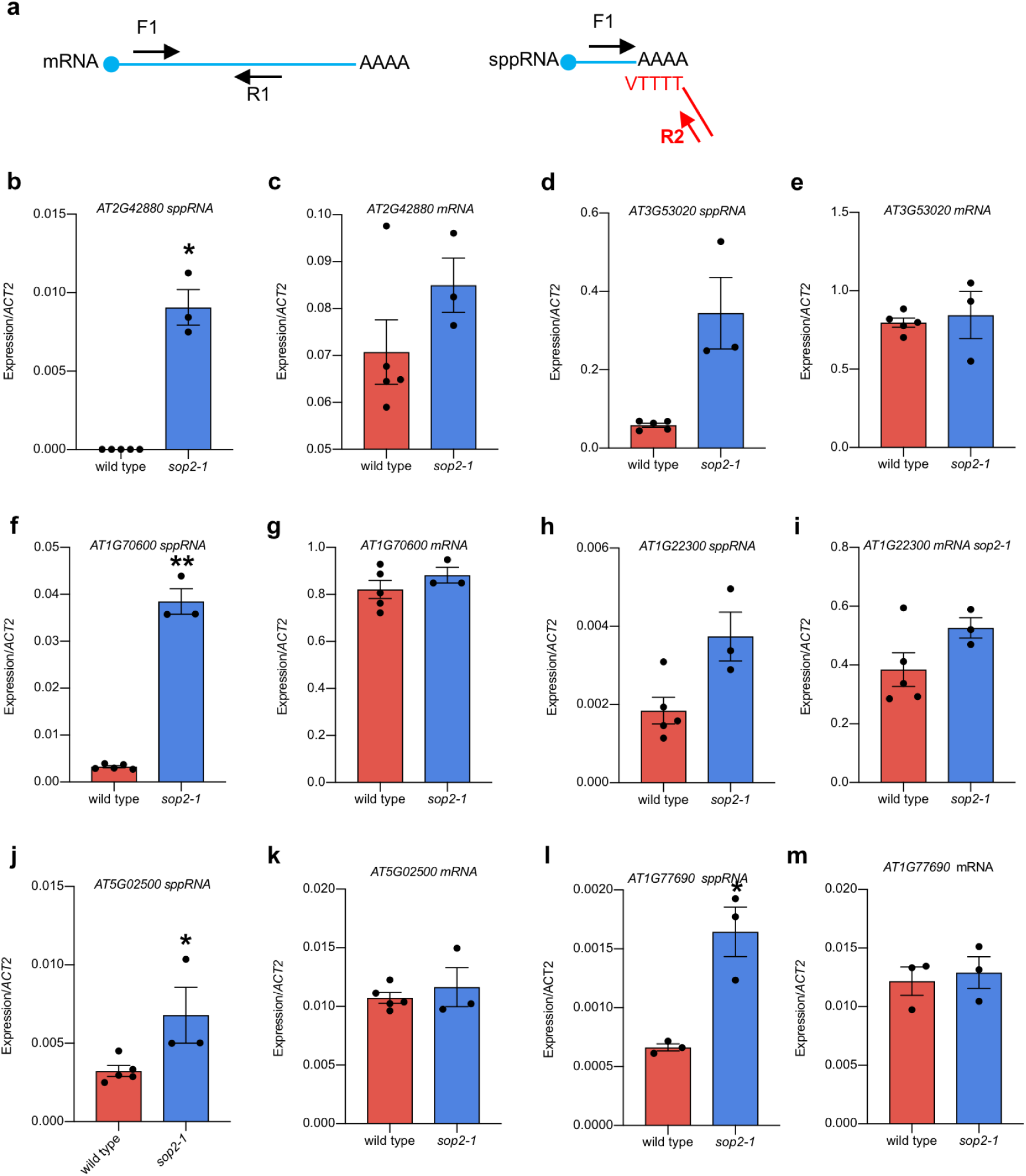
Stabilizing sppRNAs in exosome subunit RRP4 mutant *sop2-1* does not significantly change cognate mRNA expression. (a) Schematic representation of RT-qPCR-based strategy to selectively amplify sppRNAs and mRNAs using very short annealing and extension times (see Methods). RT-qPCR analysis of sppRNA levels (using F1/R2 primer pairs) and mRNA levels (using F1/R1 primer pairs) detected in wild type and *sop2-1(Hematy, Bellec et al. 2016)* for (b-c) *MPK20 (AT2G42880)*, (d-e) *RPL24 (AT3G53020)*, (f-g) *RPL18e (AT1G70600)*, (h-i) *GF14 (AT1G22300)*, (j-k) *HSC70-1 (AT5G02500)*, and (l-m) *LAX3* (*AT1G77690)*. Black dots indicate individual data points. Error bars represent SEM resulting from at least three independent experiments. For statistical test, a single asterisk denotes p<0.05, while two asterisks denotes p<0.01, between wild type and *sop2-1* by Student’s t-test.

**Figure S11.**
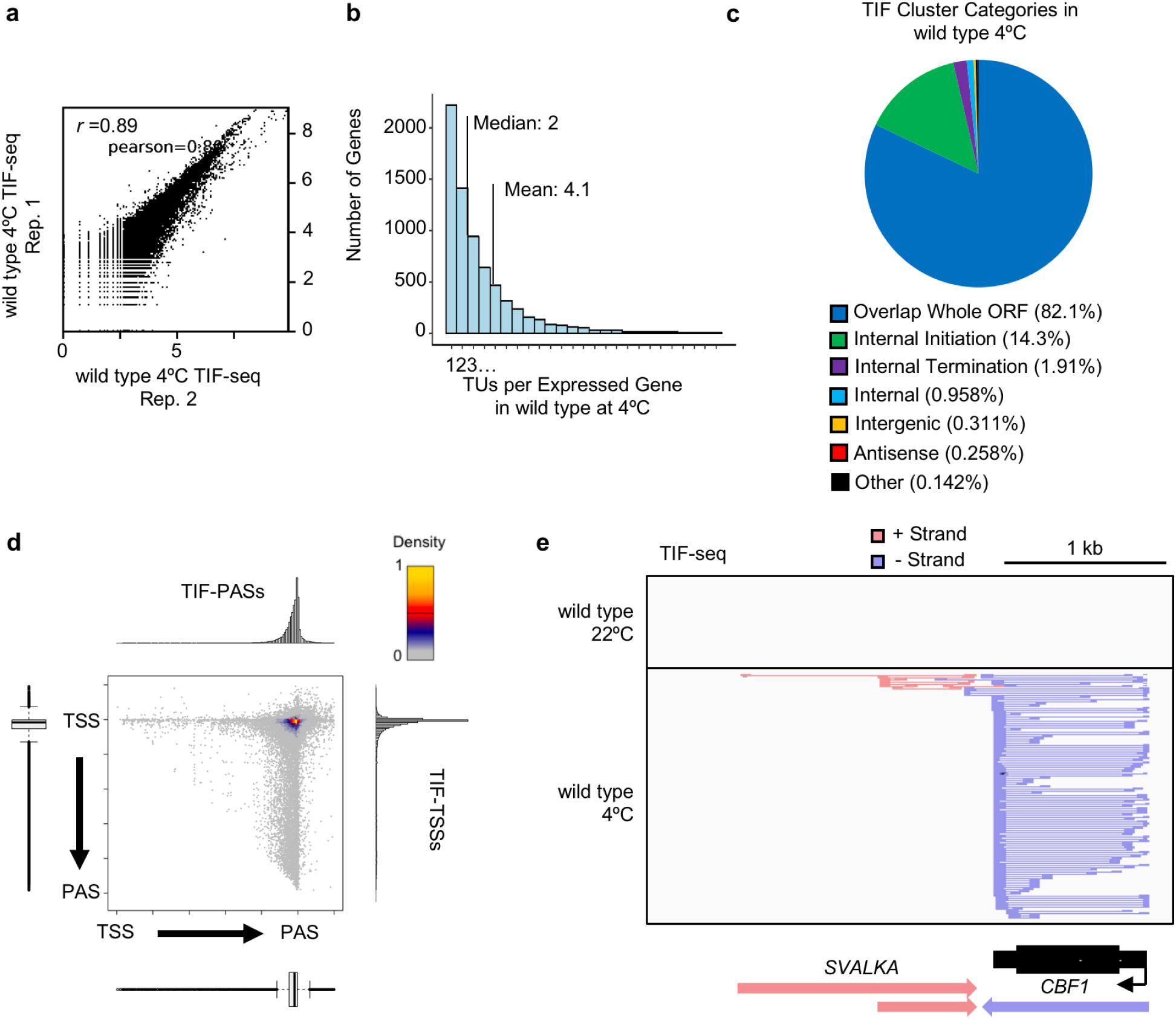
TIF-seq analyses in cold-treated wild type seedlings. (a) Scatterplot of log_10_(coverage)/5 nt bins for TIF-reads from two independent replicates in wild type seedlings following three hours at 4°C (see Methods). (b) Histogram of number of TIF-clusters per expressed gene in cold-treated wild type seedlings (mean: 4.2; median: 2). (c) Pie-chart displaying the proportion of TIF Cluster Categories detected in cold-treated wild type seedlings. (d) Scatterplot of TSS/PAS pairs detected by TIF-seq in wild type, with respect to genome annotations. Histograms and bar plots display metagene TIF-TSS and TIF-PAS distribution normalized to gene length. (e) Genome browser screenshot depicting TIF-seq detection of cold-induced *C-REPEAT BINDING FACTOR 1* (*CBF1*) gene expression and both isoforms of the recently characterized lncRNA *SVALKA* that regulate optimal *CBF1* expression and plant cold acclimation*(Kindgren, Ard et al. 2018)*.

**Figure S12.**
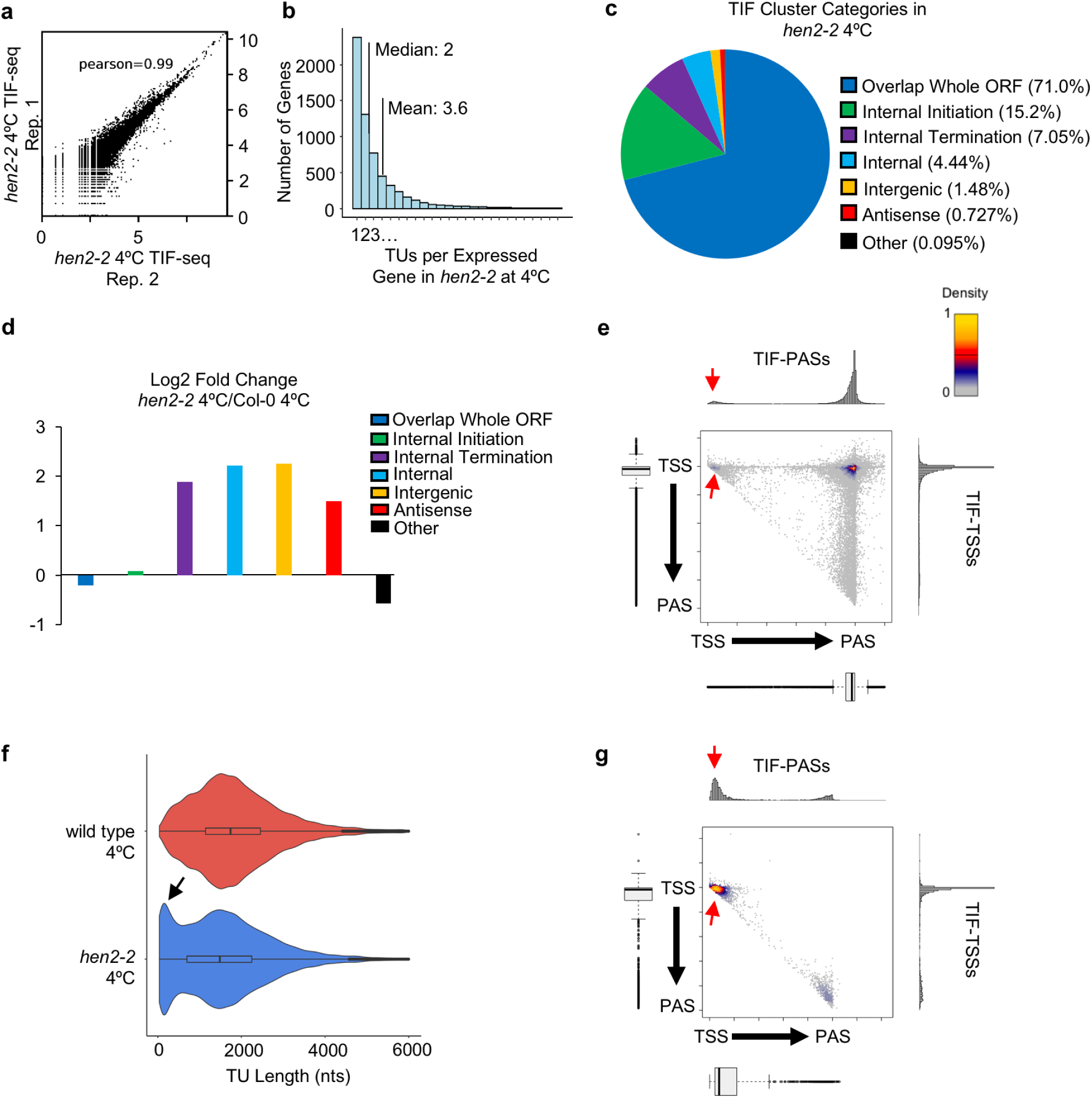
TIF-seq analyses in cold-treated *hen2-2* identify widespread premature termination and sppRNA production following environmental change. (a) Scatterplot of log_10_(coverage)/5 nt bins for TIF-reads from two independent replicates in *hen2-2* seedlings following three hours at 4°C (see Methods). (b) Histogram of number of TIF-clusters per expressed gene in cold-treated *hen2-2* seedlings (mean: 3.6; median: 2). (c) Pie-chart displaying the proportion of TIF Cluster Categories detected in cold-treated *hen2-2* seedlings. (d) log2 fold change of TIF Cluster category proportions in cold-treated *hen2-2* compared to cold-treated wild type. (e) Scatterplot of TSS/PAS pairs detected by TIF-seq in cold-treated *hen2-2*, with respect to annotated TSSs and PASs. Histograms and bar plots display metagene TIF-TSS and TIF-PAS distribution normalized to gene length. Red arrows indicate termination within initiation regions. (f) TIF-cluster sizes in cold-treated wild type and *hen2-2*. Black arrow indicates enrichment of short TUs in cold-treated *hen2-2*. (g) Scatterplot of short (<200 nt) TSS/PAS pairs detected by TIF-seq in cold-treated *hen2-2*, with respect to annotated TSSs and PASs.

**Figure S13.**
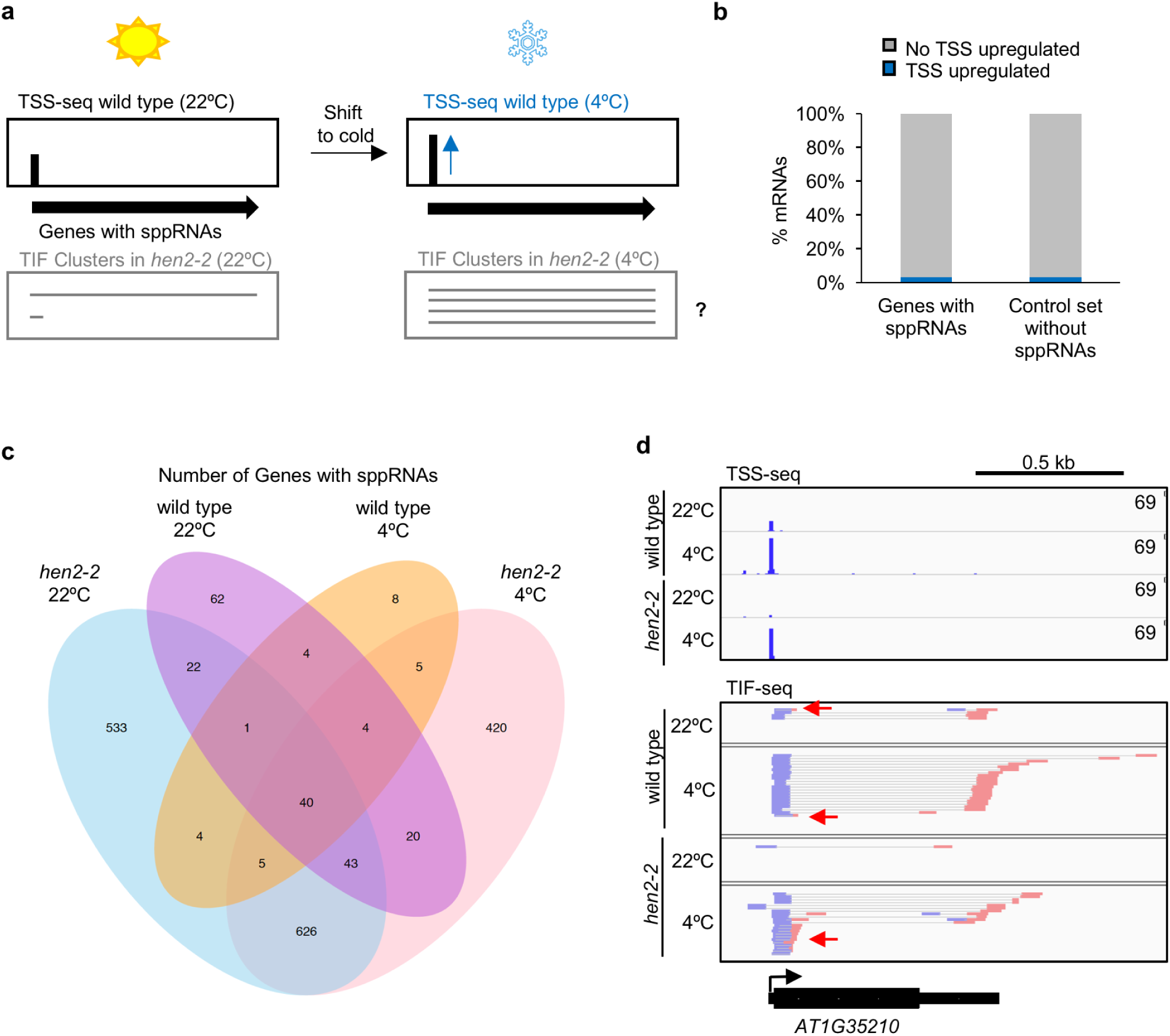
Cold treatment stimulates sppRNA production at many induced genes. (a) Illustration of possible effects of premature termination and sppRNA production on full length gene expression following environmental change. (b) Percentage of sppRNA genes that increase mRNA expression at the expense of sppRNAs following cold treatment compared to control set of genes without detectable sppRNAs. For both classes tested: n=1153. See methods. (c) Venn diagram depicting number of genes with detectable sppRNAs in wild type and/or *hen2-2* before and after 3 hours of cold treatment. See **Table S5 and S6** for sppRNA gene lists and sppRNA genomic coordinates, respectively. (d) Genome browser screenshot of TSS- and TIF-seq at the *AT1G35210* gene in wild type and *hen2-2* both before and after cold treatment. Increased *AT1G35210* mRNA levels correspond with the elevated sppRNA levels following cold treatment. The red arrow indicates sppRNAs.

**Figure S14.**
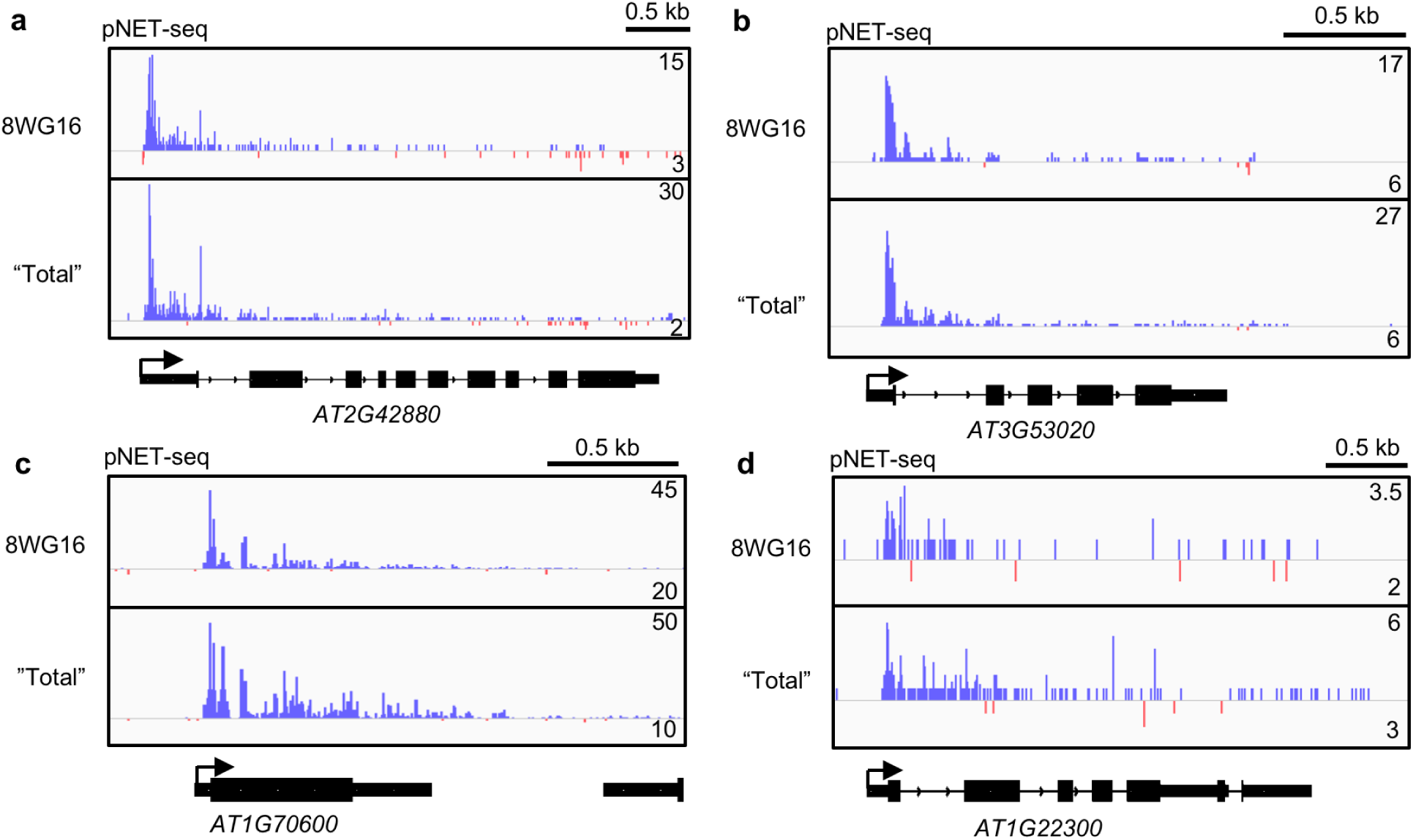
Promoter-proximal RNAPII peaks at genes with sppRNAs. Genome browser screenshots of pNET-seq data from experiments using different RNAPII antibodies, 8WG16 and “Total” RNAPII(Zhu, Liu et al. 2018), along sppRNA-containing genes: (a) *MPK20 (AT2G42880)*, (b) *RPL24 (AT3G53020)*, (c) *RPL18e (AT1G70600)*, and (d) *GF14 (AT1G22300).* Blue reads on the positive (sense) strand. Red reads on the negative (antisense) strand.

**Figure S15.**
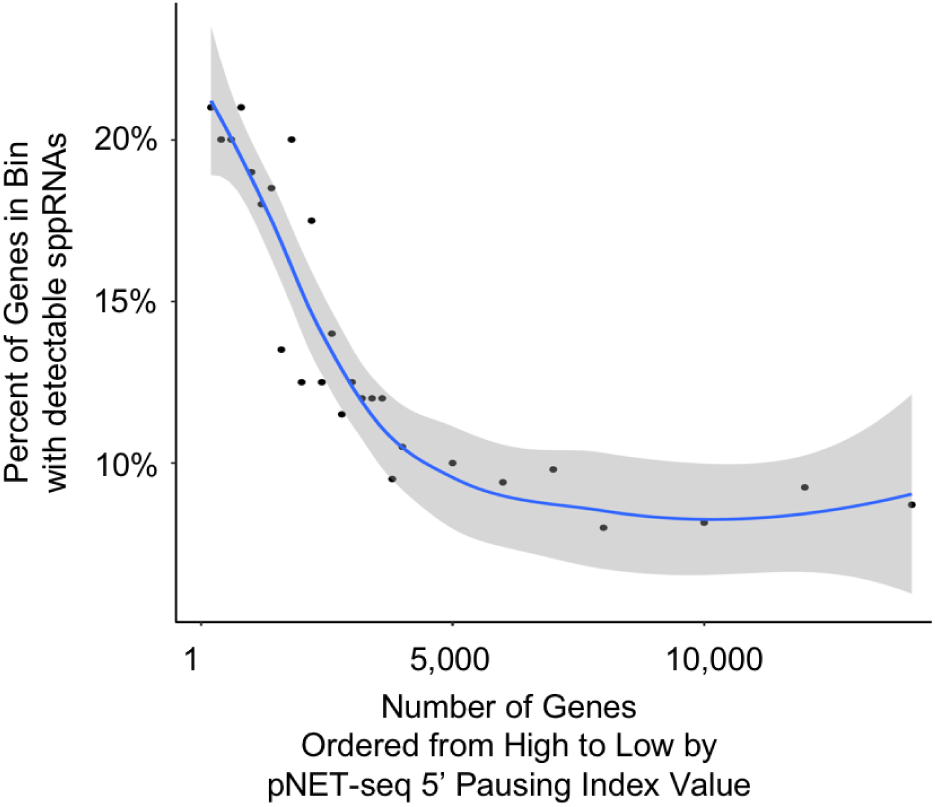
Increased promoter-proximal RNAPII stalling strongly correlates with sppRNA production. “5’ pausing index” values were calculated for 14,126 of the 75% most expressed genes from pNET-seq (8WG16)(Zhu, Liu et al. 2018) (see Methods). The percentage of genes that have detectable sppRNAs per bin of genes (beginning with 200-gene bins and ending with 2,000 gene bins) are represented as individual black dots. The trend line is indicated in blue, while shading represents a confidence interval of 95%. Overall, higher levels of promoter-proximal RNAPII stalling strongly correlates with sppRNA formation.

**Figure S16.**
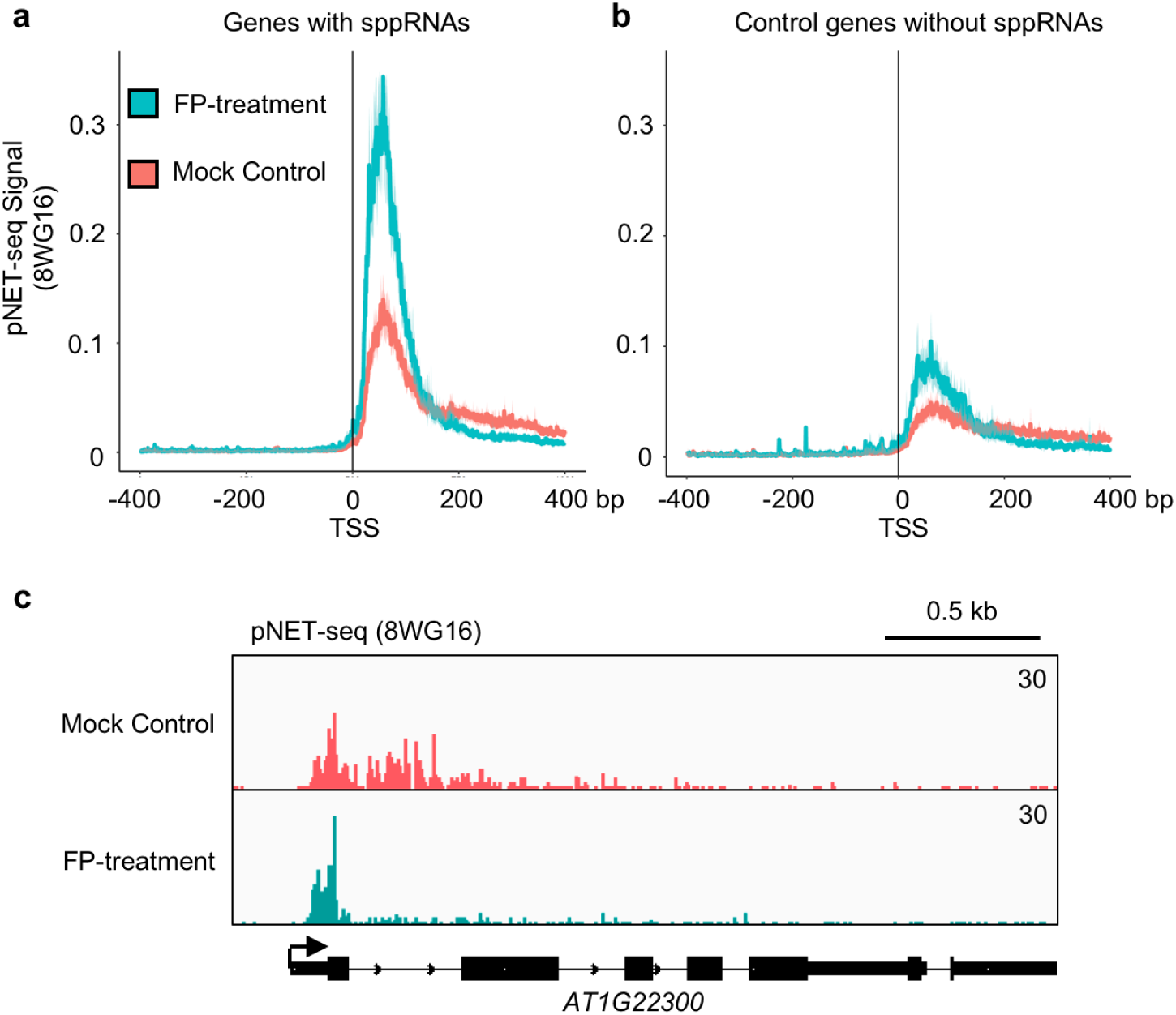
Promoter-proximal RNAPII peaks at sppRNA-producing genes are highly sensitive to PTEF-b inhibitor Flavopiridol. Metagene profiles of pNET-seq data from *Arabidopsis* seedlings before (mock) and after flavopiridol (FP) treatment(Zhu, Liu et al. 2018): (a) Genes with sppRNAs; (b) Control set of genes without detectable sppRNAs but displaying equal nascent gene body transcription as those in panel A. Gene body is defined as +200 nts from annotated TSS and - 200 nts from annotated PAS (see Methods). (c) Genome browser screenshot of pNET-seq signal along the sppRNA-containing gene *GF14 (AT1G22300)* following mock or FP treatment.

